# *Klebsiella pneumoniae* causes bacteremia using factors that mediate tissue-specific fitness and resistance to oxidative stress

**DOI:** 10.1101/2023.02.23.529827

**Authors:** Caitlyn L. Holmes, Alexis E. Wilcox, Valerie Forsyth, Sara N. Smith, Bridget S. Moricz, Lavinia V. Unverdorben, Sophia Mason, Weisheng Wu, Lili Zhao, Harry L.T. Mobley, Michael A. Bachman

## Abstract

Gram-negative bacteremia is a major cause of global morbidity involving three phases of pathogenesis: initial site infection, dissemination, and survival in the blood and filtering organs. *Klebsiella pneumoniae* is a leading cause of bacteremia and pneumonia is often the initial infection. In the lung, *K. pneumoniae* relies on many factors like capsular polysaccharide and branched chain amino acid biosynthesis for virulence and fitness. However, mechanisms directly enabling bloodstream fitness are unclear. Here, we performed transposon insertion sequencing (TnSeq) in a tail-vein injection model of bacteremia and identified 58 *K. pneumoniae* bloodstream fitness genes. These factors are diverse and represent a variety of cellular processes. *In vivo* validation revealed tissue-specific mechanisms by which distinct factors support bacteremia. ArnD, involved in Lipid A modification, was required across blood filtering organs and supported resistance to soluble splenic factors. The purine biosynthesis enzyme PurD largely enhanced liver fitness and was required for replication in serum. PdxA, a member of the endogenous vitamin B6 biosynthesis pathway, optimized replication in serum and lung fitness. The stringent response regulator SspA was required for splenic fitness yet was dispensable in the liver. In a bacteremic pneumonia model that incorporates initial site infection and dissemination, splenic fitness defects were enhanced, and DsbA, SspA, and PdxA increased fitness across bacteremia phases. SspA and PdxA enhanced *K. pnuemoniae* resistance to oxidative stress. SspA specifically resists oxidative stress produced by NADPH oxidase Nox2 in the lung, spleen, and liver, as it was a fitness factor in wild-type but not Nox2-deficient (*Cybb*^−/−^) mice. These results identify site-specific fitness factors that act during the progression of Gram-negative bacteremia. Defining *K. pneumoniae* fitness strategies across bacteremia phases could illuminate therapeutic targets that prevent infection and sepsis.

**Author Summary:** Gram-negative bacteremia is a deadly family of infections that initiate sepsis, a leading cause of global morbidity and mortality. Only a small number of Gram-negative species contribute to the majority of clinical bacteremia. *Klebsiella pneumoniae* is the second leading cause of Gram-negative bacteremia, and the third leading cause of overall bloodstream infection. *K. pneumoniae* is highly linked to hospital-associated infection with increasing antimicrobial resistance, endangering the most vulnerable patients. It is critical to understand the pathogenesis of *K. pneumoniae* bacteremia to better develop targets for future therapies that can prevent these deadly infections. Here, we define over 50 *K. pneumoniae* genes that support bloodstream fitness. These factors are diverse, support tissue-specific fitness, and increase bacterial resistance to oxidative stress. Our study is the first to systematically define *K. pneumoniae* factors enhancing bacteremia in a mammalian system. These results illuminate host-pathogen interactions during *K. pneumoniae* bacteremia that may be extended to additional Gram-negative species.

## Introduction

Bacteremia, the presence of bacteria in the bloodstream, can initiate sepsis. Defined as immune dysregulation resulting in organ dysfunction, sepsis is a significant cause of global morbidity and mortality (1, 2, 3). Gram-negative species underlie about half of clinical bacteremia cases and are emerging in dominance (4). To establish bacteremia, Gram-negative species follow three phases of pathogenesis. First, bacteria invade or colonize tissues that serve as initial sites of infection. Second, bacteria cross host barriers unique to the initial site and disseminate into the blood. Third, bacteria must survive in the bloodstream by exercising metabolic flexibility and avoiding clearance in filtering organs like the spleen and liver (2). Defining bacterial factors that enhance bacteremia is a step toward treating this family of deadly infections.

*Klebsiella pneumoniae* is the second leading cause of Gram-negative bacteremia (4). Repeatedly classified as a pathogen of urgent concern by the World Health Organization (5, 6), infection with *K. pneumoniae* is particularly problematic due to a high association with mortality and antimicrobial resistance (7, 8). *K. pneumoniae* is highly linked to hospital associated infection, particularly pneumonia (9). Accordingly, *K. pneumoniae* lung fitness mechanisms have been extensively described and include a wide variety of factors including capsular polysaccharide, branched chain amino acid synthesis, and production of citrate synthetase (10, 11). *K. pneumoniae* mechanisms enhancing dissemination have been less thoroughly described but are likely partially dependent on ADP-heptose biosynthesis and siderophores (12, 13). The host hypoxia-inducible factor 1α (HIF-1α) in lung epithelial cells also promotes dissemination, although interactions at this step are not well understood (14).

*K. pneumoniae* factors involved in the last phase of bacteremia, survival in the blood and filtering organs, remain incompletely defined. *In vitro* studies using human serum identified *K. pneumoniae* capsule biosynthesis and lipopolysaccharide (LPS) O-antigen as essential for complement resistance (15, 16). Human serum studies also demonstrated that vitamin biosynthesis and protein translocation using the *tat* system support maximum growth (16). Modeling bacteremia using *Galleria mellonella* further confirmed a role for capsular polysaccharide, LPS, cellular envelope integrity, and iron acquisition systems during infection (17). While *in vitro* studies have consistently identified subsets of *K. pneumoniae* genes required for serum growth and complement evasion, these models cannot define factors that influence fitness in blood filtering organs. Since bloodstream fitness involves both replication and evasion of clearance, some factors may be dispensable for serum growth but perpetuate bacteremia through interactions within tissues. For example, GmhB, an ADP-heptose biosynthesis enzyme required to produce intact LPS inner core, is dispensable for serum growth and lung fitness but is required for fitness in the liver and spleen during bacteremia (12). Thus, GmhB is an example of a fitness factor specific to the last phase of bacteremia and can only be observed *in vivo*. This demonstrates that approaches defining bloodstream survival mechanisms should incorporate systemic infection to fully reveal *K. pneumoniae* factors perpetuating bacteremia.

Additionally, *in vivo* models can uncover host-pathogen interactions during bacteremia. To eliminate bacteria, immune cells may induce an oxidative stress response. NADPH oxidases, specifically the phagocytic Nox2, generate bursts of reactive oxygen species (ROS) that are frontline host defense mechanisms during infection (18). Despite the importance of this response, *K. pneumoniae* factors enhancing resistance to ROS-mediated stress have not been detected in blood filtering organs. Immune responses can also vary across organs as tissue-resident cells can have differential interactions with bacteria, including *K. pneumoniae* (19). Varying replication rates of multiple Gram-negative species across tissues during bacteremia highlights that host-pathogen interactions are not uniform between sites (20). Thus, defining bacterial site-specific fitness mechanisms can illuminate host defense strategies.

Our group has used transposon insertion-site sequencing (TnSeq) to define bacterial genes required for bloodstream fitness in Gram-negative species including *Escherichia coli, Serratia marcescens, Citrobacter freundii, Acinetobacter baumannii*, and *Proteus mirabilis* (21, 22, 23, 24, 25, 26). These studies defined an array of bacterial genes enhancing bloodstream fitness. Metabolic flexibility emerged as a prominent feature by which Gram-negative species survive in the blood. Interestingly, no factors have been identified that are universally required across all six species. This highlights that bacteremia may require a species-specific arsenal of fitness mechanisms. Considering the dominance of *K. pneumoniae* infections in clinical settings and resulting high mortality rates (7, 8, 9, 27), it is critical to define mechanisms by which this specific pathogen perpetuates bacteremia. Additionally, *K. pneumoniae* mechanisms of pathogenesis apart from the lung are largely unknown and gaining knowledge of these processes will allow insight into this pathogen.

To define *K. pneumoniae* bacteremia factors influencing bloodstream survival, we performed TnSeq using a mammalian model of intravascular bacteremia and revealed 58 diverse genes that enhance splenic fitness. Validation studies demonstrate that bacteremia is enhanced by a set of factors relaying tissue-specific fitness in the spleen, serum, and liver. This study is the first to systematically identify *K. pneumoniae* strategies for *in vivo* bloodstream fitness.

## Results

### Transposon insertion-site sequencing during *K. pneumoniae* bacteremia reveals diverse mechanisms enhancing bloodstream fitness

To define *K. pneumoniae* factors influencing bacteremia pathogenesis, transposon-insertion site sequencing (TnSeq) was performed using a previously described KPPR1 library containing ~25,000 unique mutations (11, 28). To model the third phase of bacteremia, survival in the blood and filtering organs, an intravascular model was used with inoculation of mice via tail vein injection (Fig 1A). This model bypasses the first two phases of bacteremia, initial site infection and dissemination, and allows for direct examination of bloodstream fitness (12, 29). To define potential experimental bottlenecks to the spleen in this model, a fitness-neutral mutant was competed against wild-type KPPR1 at ratios of 1:1, 1:5,000, and 1:10,000. At the 1:10,000 ratio, the mutant was recovered in significantly lower abundance than KPPR1 indicating potential stochastic loss at this ratio (S1 Fig). In contrast, loss was minimal at the 1:5,000 dilution. Therefore, the splenic bottleneck after tail vein injection was determined to be between these dilutions, and 1:8,500 was selected as the desired maximum input complexity to represent approximately 1/3 of the total mutants. To increase the chance of sampling all mutants in the library, the KPPR1 transposon library was used to generate four pools with ~8,500 unique insertions each, and each pool (Pools A-D) was administered to 10 mice.

**Fig 1.**
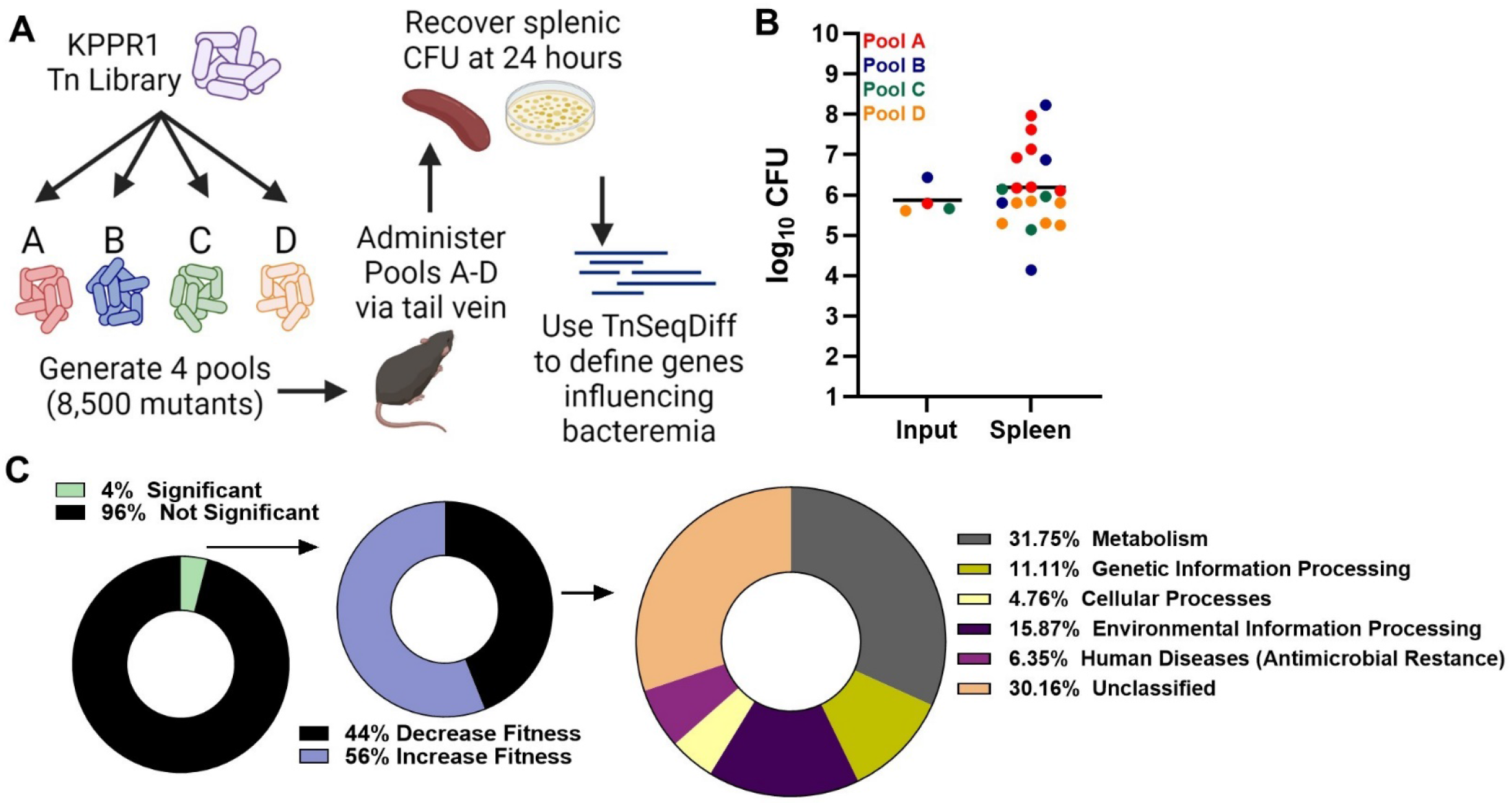
Transposon insertion-site sequencing (TnSeq) reveals that *K. pneumoniae* bacteremia is enhanced by a set of genes representing diverse fitness mechanisms. (A) Overview of *K. pneumoniae* bacteremia TnSeq. A KPPR1 transposon library was divided into four pools containing 8,500 unique insertions and administered to mice via the tail vein at a 1×10^6^ CFU dose. After 24 hours, splenic CFU was recovered, and DNA was sequenced. The TnSeqDiff pipeline determined genes influencing fitness. (B) The input and splenic CFU burden for each pool and mouse represented in TnSeq. (C) Genes represented in TnSeq (~3,800 genes) that were defined as influencing infection (132 genes), and primary KEGG orthology for genes increasing *K. pneumoniae* fitness during bacteremia (58 genes).

Twenty-four hours post-inoculation, total splenic CFU was recovered, DNA was extracted and subjected to sequencing, and the abundance of unique transposon insertions was compared to the inoculum as described in the Materials and Method section. Mean splenic colonization was roughly 1×10^6^ CFU (Fig 1B) and mouse mortality between pools varied widely from 0-70%. As a result, spleen samples from 10, 3, 9, and 10 mice administered pools A, B, C, and D, respectively, were available for analysis. Within the input pools, ~3,800 of the estimated 4,312 non-essential KPPR1 genes contained transposon insertions (S1 Table). Transposon mutations within individual genes varied by pool, ranging from 9,350-12,450 unique transposon insertions. 148 genes, 4% of those within the study, were identified as significantly influencing *K. pneumoniae* fitness during bacteremia (Fig 1C). Of significant hits, 58 (44%) genes with transposon insertions resulted in lower recovery compared to the input, revealing bacterial factors enhancing bacteremia. KEGG annotation summaries for these genes demonstrated that a diverse set of *K. pneumoniae* factors enhance bloodstream survival (Table 1). The most highly represented genetic function was metabolism (Fig 1C). Defined by KEGG orthology, no single metabolism type was dominant and synthesis of carbohydrates, nucleotides, amino acids, vitamins, and other substrates supported bloodstream fitness (S2 Fig). In addition, 56% of genes with transposon insertions resulted in higher recovery compared to the input (S2 Table). This pattern indicates a set of genes that suppress pathogenesis since mutations led to enrichment after infection. Most of these hits have unclassified functional KEGG annotations. Of the hits with annotated functions, metabolic pathways were also highly represented in this group (S2 Fig). Since the goal of the present study was to define genes enhancing bacteremia, this group was not analyzed further.

**Table 1.**
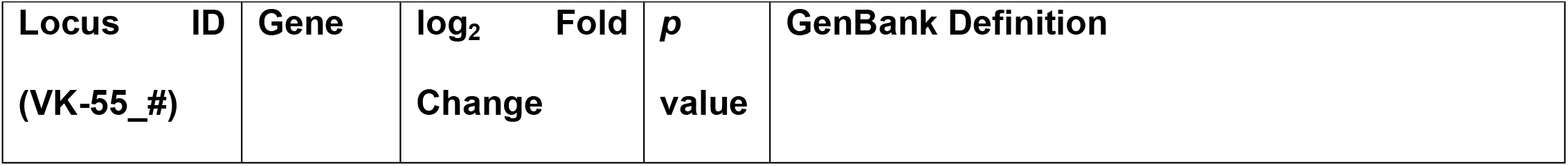
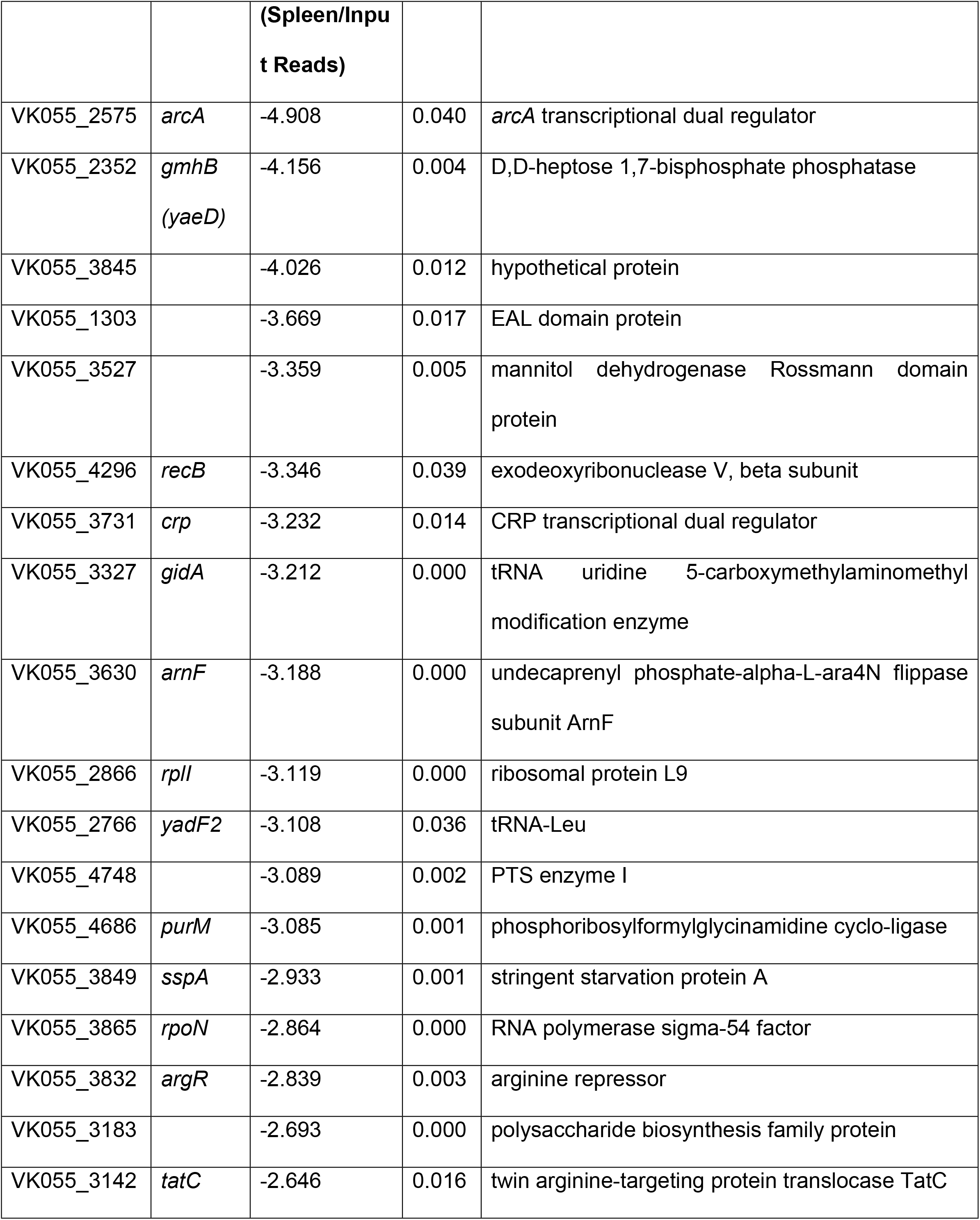
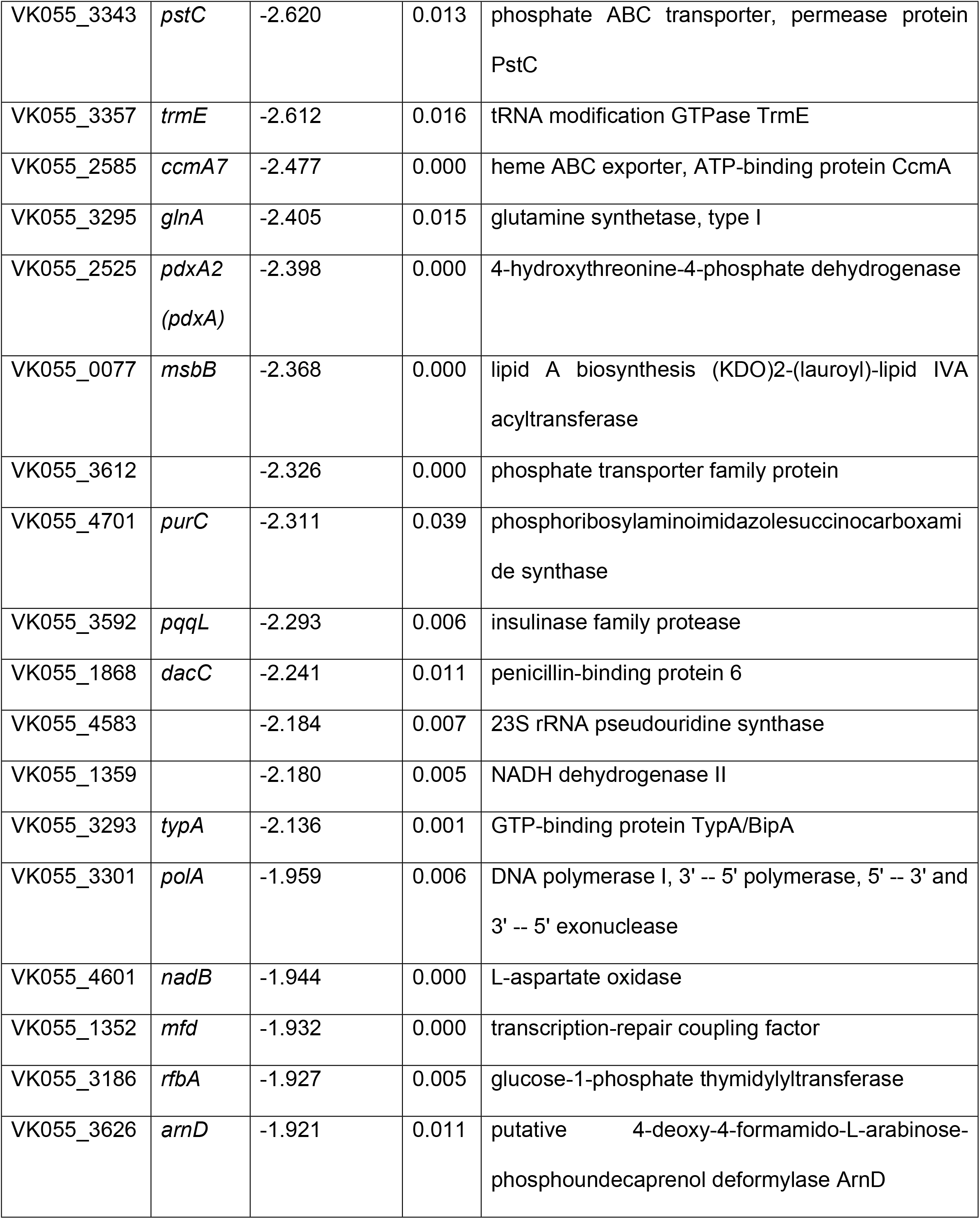
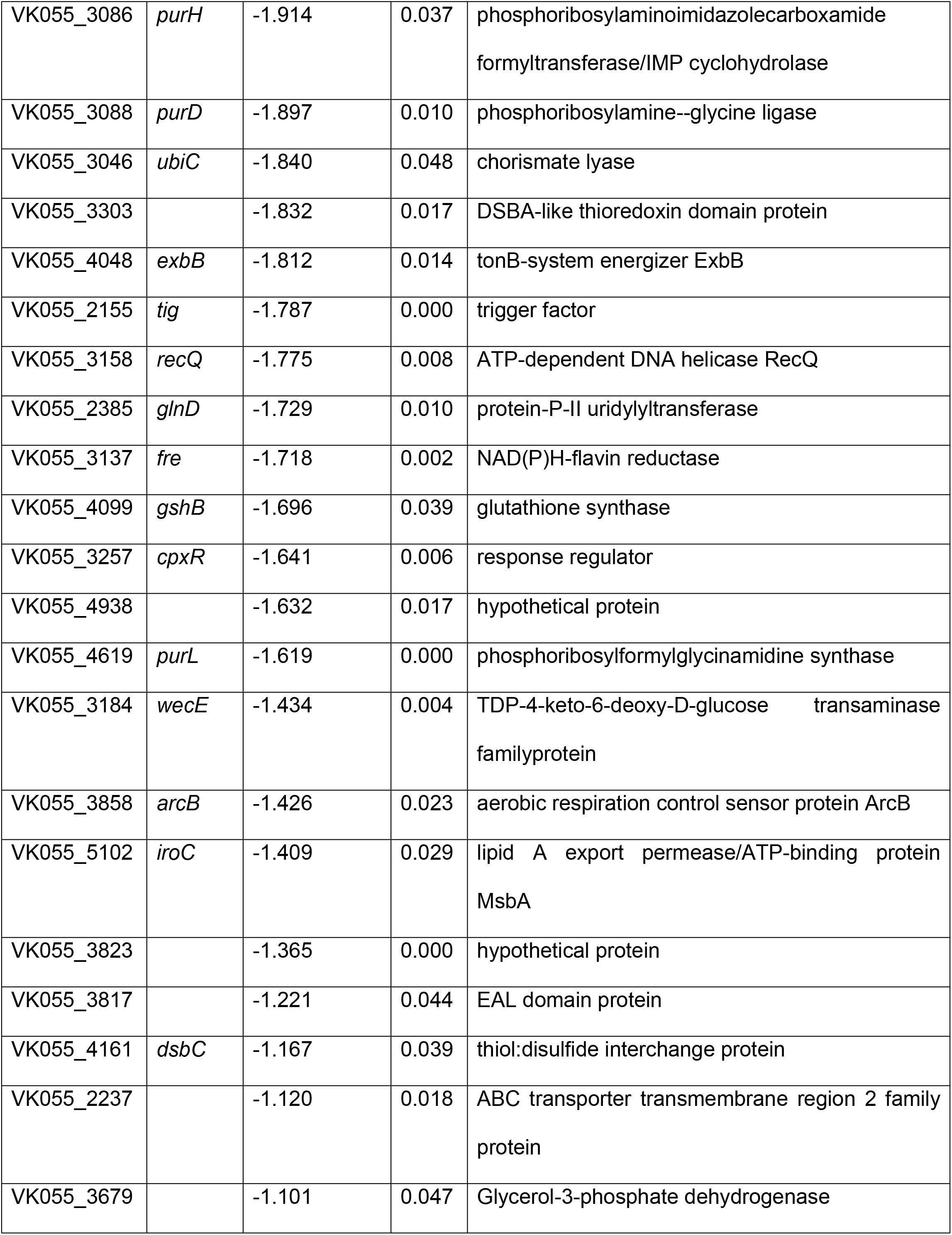
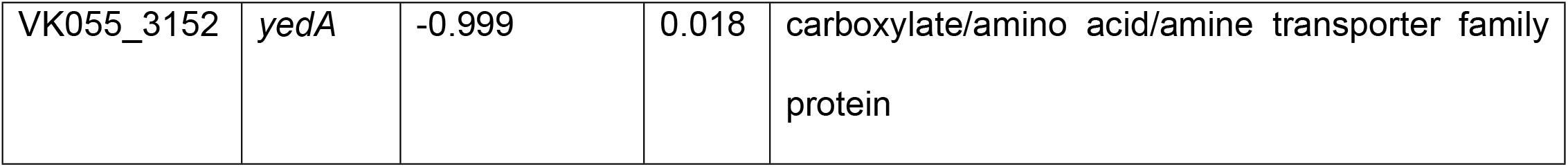
*K. pneumoniae* splenic fitness factors.

Six of the 58 genes enhancing bacteremia were then selected for characterization based on representing diverse cellular functions, potential conserved Enterobacterales bacteremia fitness factors (22, 23), or being unique to *K. pneumoniae*. Transposon mutants for each gene of interest were selected from a KPPR1 ordered library (30) and growth was observed in rich medium (LB). Apart from *glnA*, a glutamine synthetase, *in vitro* replication was not influenced by mutations within these genes, meaning that contributions to bacteremia were likely unrelated to basic cellular replication in nutrient rich environments (S3 Fig). *K. pneumoniae* bacteremia hits also had varying effects on hypermucoviscosity (S3 Fig). Mutations within *arnD* significantly reduced, as previously reported (30), while mutations within *glnA* and *pdxA* enhanced, hypermucoviscosity.

### *K. pneumoniae* bacteremia pathogenesis is perpetuated by factors that relay site-specific fitness

To validate the bacteremia TnSeq results, the five genes of interest that did not demonstrate *in vitro* replication defects were further analyzed: *arnD, purD, dsbA, sspA*, and *pdxA*. Each gene was represented by 3-5 unique transposon insertions within the TnSeq study, and insertion level analysis for each gene’s unique mutations across individual animals demonstrated consistent loss in splenic abundance compared to the input (S4 Fig).

For *in vivo* validation of the TnSeq study, individual transposon mutants were competed against KPPR1 at a 1:1 ratio in the intravascular bacteremia model. ArnD, a member of a Lipid A modification system, had the greatest influence on bacteremia, demonstrating an *in vivo* fitness requirement in both the spleen and liver (Fig 2A, S5 Fig). A mutant in *purD*, involved in endogenous purine biosynthesis, approached but did not reach significance (*P*=0.08) for splenic fitness defects, but was significantly defective in the liver (Fig 2B, S5 Fig). DsbA, a member of a disulfide bond formation and secretion system, was required in both compartments but had a larger influence on liver fitness (Fig 2C, S5 Fig). In contrast, SspA, a regulator of the stringent starvation response, was required in the spleen but was dispensable in the liver (Fig 2D, S5 Fig). PdxA, a member of the vitamin B6 biosynthesis cascade, had a modest but significant fitness defect in the spleen (Fig 2E, S5 Fig). Notably, each bacteremia factor enhanced fitness in a distinct manner across blood filtering organs. Factors displayed unique patterns of tissue-specific fitness with some, like ArnD and DsbA being required across organs, while PurD, SspA, and PdxA were only required in one organ.

**Fig 2.**
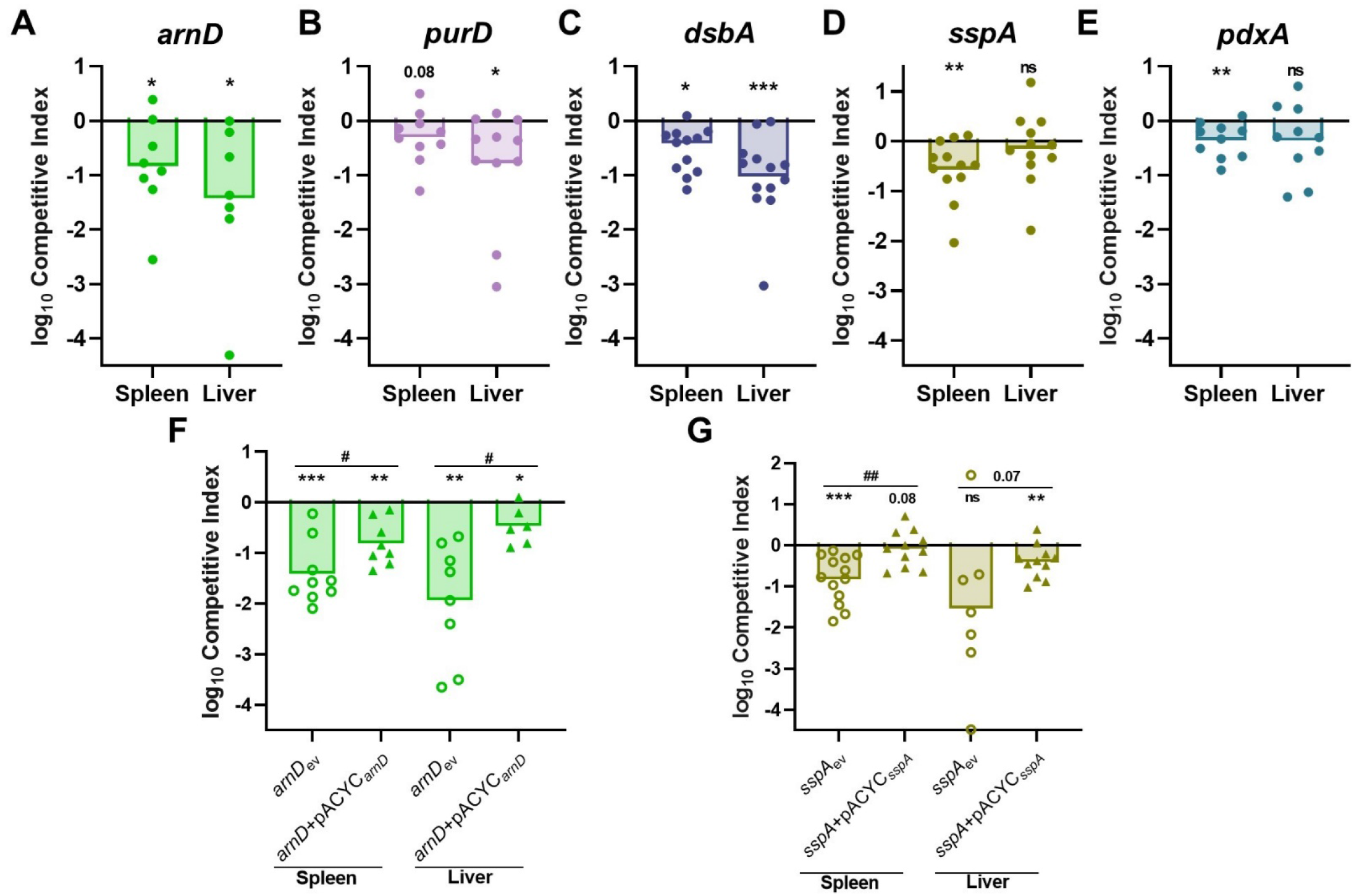
*K. pneumoniae* bacteremia fitness factors directly relay tissue-specific fitness advantages. Five factors indicated by TnSeq as significantly enhancing bacteremia were selected for *in vivo* validation using the tail vein injection model. The 1:1 inoculum consisted of KPPR1 and transposon mutants for (A) *arnD*, (B) *purD*, (C) *dsbA*, (D) *sspA*, or (E) *pdxA*. Competitions were also performed using strains carrying the empty pACYC vector (_ev_) or complementation provided on pACYC184 under native promoter control for (F) *arnD* (*arnD*+pACYC_*arnD*_) or (G) *sspA* (*sspA*+pACYC_*sspA*_). Mean log_10_ competitive index at 24 hours post infection is displayed. For all, *p<0.05, **p<0.01, ***p<0.001 by one sample t test with a hypothetical value of zero; for (F, G) ^#^p<0.05, ^##^p<0.01 by unpaired *t* test. For each group, n*≥*8 mice in at least two independent trials.

To confirm that fitness defects were specifically related to disruption of the genes of interest, *arnD* and *sspA* were complemented *in trans* as mutations in these genes resulted in the largest splenic fitness defects. Complementation of *arnD* significantly alleviated bacteremia fitness defects in comparison to *arnD*_ev_, demonstrating that *arnD* enhances splenic and liver fitness (Fig 2F, S5 Fig). However, the *arnD+*pACYC_*arnD*_ strain continued to demonstrate lower fitness in relation to KPPR1_ev_. This is likely due to polar effects on downstream genes of the collective function of the *arn* system, consisting of other Lipid A modifying enzymes (31). Indeed, another member of the *arn* operon, *arnF*, was a TnSeq hit, further validating a role for this system in bacteremia (Table 1). SspA is a regulator of the complex stringent starvation response and *sspA* mutations lead to higher susceptibility to many environmental stressors (32). SspA complementation (*sspA+*pACYC_*sspA*_) significantly ameliorated *K. pneumoniae* splenic fitness defects (Fig 2G, S5 Fig). Therefore, *in vivo* complementation of both selected factors restored *K. pneumoniae* bacteremia fitness and specifically highlight genes that influence spleen and liver fitness. This confirms that the TnSeq study revealed multiple factors that directly enhance bacteremia as site-specific factors.

### *K. pneumoniae* utilizes multiple strategies to enhance site-specific fitness during bacteremia, including metabolic flexibility, serum resistance, and LPS modification

Since *in vivo* validation revealed distinct fitness factor contributions to tissue-specific fitness, individual bloodstream compartments were investigated to determine relevant interactions during infection. Bacterial replication occurs across compartments during bacteremia, and metabolic flexibility is critical to growth in the bloodstream (20, 21, 22). To test contributions to growth in serum, each of the five validated bacteremia fitness factors were assessed for *in vitro* replication in mouse serum (Fig 3, S6 Fig). Purine biosynthesis, mediated by PurD, was required for growth in murine and human serum as previously described (17, 33). This defect was not explained by complement resistance, as the *purD* mutant had a similar growth defect in both active and heat inactivated serum (Fig 3B-C, S6 Fig). Genetic complementation of *purD* restored the ability for serum growth compared to the wild-type or *purD* strains carrying an empty complementation vector (Fig 3D, S6 Fig). Purines were sufficient to restore serum growth as exogenous purine supplementation ameliorated *purD* defects (Fig 3E, S6 Fig). Vitamin B6 biosynthesis also maximized replication in murine and human serum, as a *pdxA* mutant showed a mild defect (Fig 3A-C). Genetic complementation of *pdxA* restored normal replication compared to wild-type and *pdxA* strains carrying the empty complementation vector (Fig 3F, S6 Fig). This indicates that the serum is a nutrient restricted environment and *K. pneumoniae* must endogenously produce key metabolites including purines and vitamin B6 to enable bloodstream replication.

**Fig 3.**
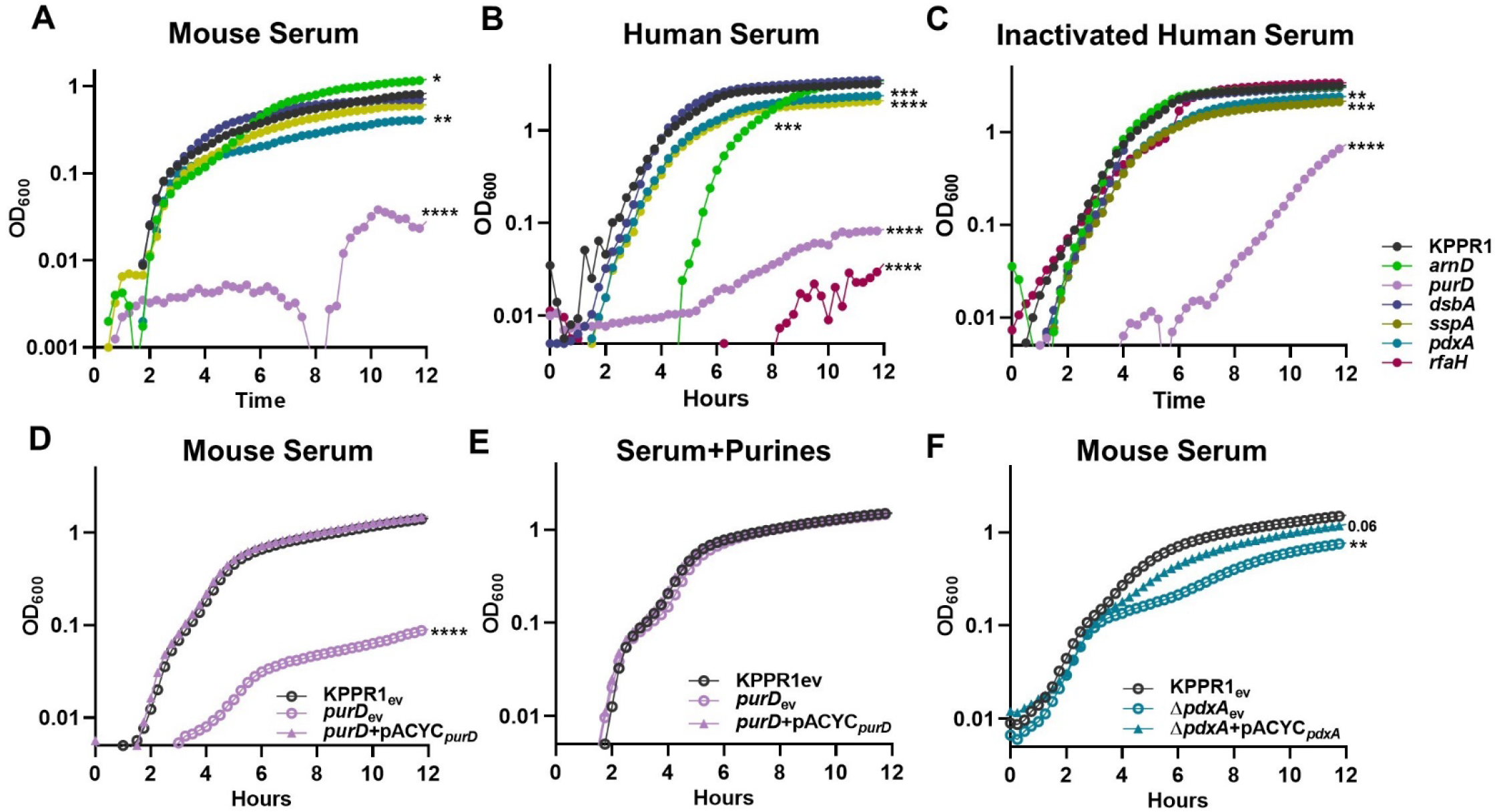
*K. pneumoniae* serum replication requires purine synthesis and is maximized by endogenous vitamin B6 biosynthesis. *K. pneumoniae* strains were grown in M9 salts supplemented with (A) 10% mouse serum or 20% human serum that was either (B) active or (C) heat inactivated. *K. pneumoniae* strains carrying the empty vector pACYC (_ev_) or pACYC expressing (D,E) *purD* (*purD*+pACYC_*purD*_) or (F) *pdxA* (Δ*pdxA*+pACYC_*pdxA*_) were grown in 10% mouse serum (D, F). Chemical complementation for *purD* was measured by supplementation of 1mM purines prior to growth (E). For all, the OD_600_ was measured every 15 minutes for 12 hours. Differences in growth compared to KPPR1 or KPPR1_ev_ were detected by area under the curve using a one-way ANOVA with Dunnett’s multiple comparison for each strain compared to wild-type; *p<0.05, **p<0.01, ***p<0.001, ****p<0.0001. For each group, n=3.

ArnD is a member of a well-described LPS modification system (*arn* operon) that covalently attaches arabinose residues onto Lipid A (31). Mutations in *arnD* render *K. pneumoniae* significantly less hypermucoviscous and reduce capsular polysaccharide production (S3 Fig, (30)). In active human serum, the *arnD* mutant was defective for growth, which was attributable to complement-mediated killing as heat inactivation of human serum restored normal *arnD* replication (Fig 3B-C, S6 Fig). A similar pattern was observed for the control strain *rfaH*, which lacks capsular polysaccharide and is more susceptible to killing by active human serum (11, 12). Interestingly, murine serum replication was enhanced in the absence of ArnD (Fig 3A, S6 Fig). Since ArnD was dispensable for murine serum replication yet required for *in vivo* fitness in blood filtering organs, its fitness contribution in the spleen and liver was investigated (Fig 4). Using spleen and liver organ homogenates from uninfected mice, *ex vivo* competitions were performed for KPPR1 and *arnD*. Both experienced growth in organ homogenates (S7 Fig). The *arnD* mutant had a subtle yet significant fitness defect in liver homogenate and a dramatic fitness defect in splenic homogenate (Fig 4A). To define relevant splenic compartments, splenocytes were removed from the homogenate, leaving only the soluble fraction. Strikingly, the mutant was also defective in splenic filtrate. This indicates that ArnD increases protection against a soluble factor specifically found in the spleen.

**Fig 4.**
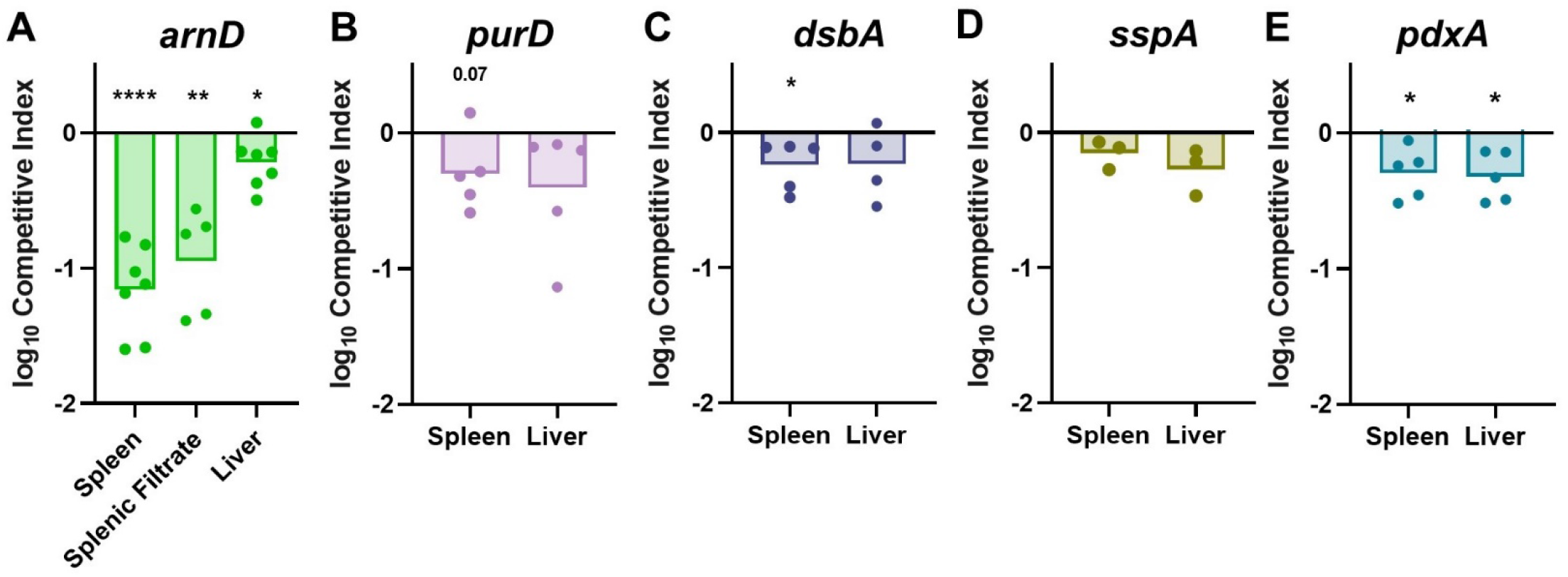
*K. pneumoniae* bacteremia factors convey fitness advantages through tissue-specific interactions within blood-filtering organs. *Ex vivo* competitions were performed using uninfected murine spleen or liver homogenate with a 1:1 mixture of KPPR1 and transposon mutants for (A) *arnD*, (B) *purD*, (C) *dsbA*, (D) *sspA*, (E) or *pdxA*. Mean log_10_ competitive index compared to wild-type KPPR1 at 3 hours post incubation is displayed. *p<0.05, **p<0.01, ****p<0.0001 by one-sample *t* test with a hypothetical value of zero and n=3-7 with points representing individual mice.

### Splenic fitness is influenced by route of infection

While ArnD and PurD dramatically influenced spleen and serum fitness, respectively, the contributions of DsbA, SspA, and PdxA to splenic fitness were more subtle (Fig 2). DsbA and PdxA were directly linked to enhancing splenic fitness as *K. pneumoniae* fitness defects in *dsbA* and *pdxA* were reproducible in uninfected splenic homogenate, mirroring the finding during infection (Fig 4C, E). PurD and SspA fitness defects in the spleen were not observed in *ex vivo* assays (Fig 4B, D). This could be due to direct, intravascular bacteremia only representing the third phase of pathogenesis and not encompassing the entire infection progression. To determine if DsbA, SspA, and PdxA contributions to splenic fitness could be further resolved, *in vivo* competitions were repeated using a bacteremic pneumonia model. As in the intravascular model, each factor promoted bacteremic pneumonia in a distinct manner. DsbA substantially supported lung fitness (Fig 5A, S8 Fig), linking DsbA to initial site fitness during bacteremia. SspA promoted fitness across sites at similar magnitudes (Fig 5B, S8 Fig), yet splenic fitness defects were more pronounced when incorporating a model with initial site infection and dissemination. Thus, utilization of the bacteremic pneumonia model allows for greater resolution of splenic fitness defects and highlights that some factors, like DsbA, may be most relevant to early phases of bacteremia.

**Fig 5.**
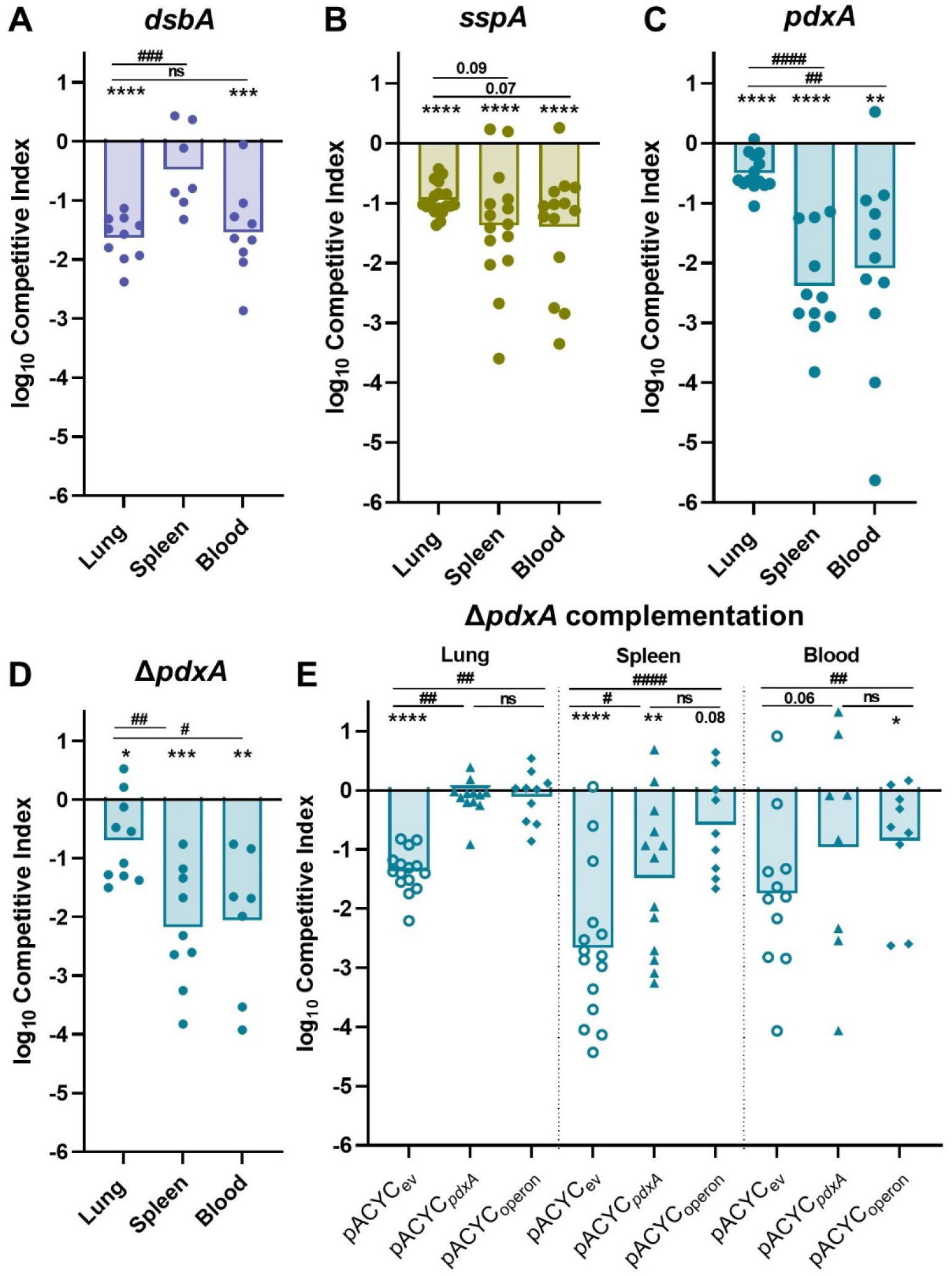
Primary site initiation of bacteremia enhances resolution of splenic fitness defects and illuminates requirements of factors across phases of pathogenesis. To model bacteremic pneumonia, mice were infected with 1×10^6^ CFU *K. pneumoniae*. Competitive infections were performed with a 1:1 mixture of KPPR1 and a transposon mutant for (A) *dsbA*, (B) *sspA*, or (C) *pdxA* or (D) a *pdxA* knockout (Δ*pdxA*). (E) Competitions were also performed with strains carrying the empty pACYC vector (_ev_) or complementation provided on pACYC for *pdxA* only (pACYC_*pdxA*_) or *pdxA* and downstream members of the operon (pACYC_operon_). Mean log_10_ competitive index at 24 hours post infection is displayed. For all, *p<0.05, **p<0.01, ***p<0.001, ****p<0.0001 by one-sample *t* test with a hypothetical value of zero; ^#^p<0.05, ^##^p<0.01, ^###^p<0.001, ^####^p<0.0001 by unpaired *t* test. For each group, n≥10 mice in at least two independent infections.

PdxA significantly enhanced initial site fitness in the lung but had a striking effect on spleen and blood fitness during bacteremic pneumonia. This pattern indicates potential contributions across all three phases of bacteremia as dissemination defects may worsen fitness at secondary sites when compared to initial sites (12). Since the splenic fitness defect was most apparent in the bacteremic pneumonia model, *pdxA* complementation was performed in this infection. However, *in trans* complementation with *pdxA* and its upstream region did not restore fitness at any site (S8 Fig). *pdxA* is a member of a large, multifunctional operon and transposon insertions within *pdxA* may have polar mutations on downstream genes. To investigate the role of this operon in bacteremia, lambda red mutagenesis was used to generate a marked Δ*pdxA* strain which demonstrated similar fitness defects as the transposon *pdxA* mutant (Fig 5D, S8 Fig). Next, the Δ*pdxA* strain was complemented with *pdxA* alone (Δ*pdxA*+pACYC_*pdxA*_) or *pdxA* plus the downstream operon genes *ksgA, apaG* and *apaH* (Δ*pdxA*+pACYC_operon_). Complementation of *pdxA* alone was sufficient to restore lung fitness completely, and significantly improved fitness in the spleen but not the blood (Fig 5E, S8 Fig). Complementation of *pdxA* and downstream genes also improved fitness across sites (Fig 5E, S8 Fig), but did not significantly alleviate defects in comparison to complementation with *pdxA* only. This indicates that complementation of *pdxA* partially restores fitness defects even when downstream genes are included. Therefore, PdxA is a fitness factor in the lung and spleen.

### Splenic fitness is enhanced by oxidative stress resistance mechanisms

Genes enhancing oxidative stress were widely represented in the significant TnSeq hits. For example, previous studies have demonstrated that mutations in *mtlD, arcA, recQ*, and *cpxR* increase susceptibility to hydrogen peroxide killing across multiple species (34, 35, 36, 37). Accordingly, survival of the *K. pneumoniae* mutants of interest after exposure to hydrogen peroxide was measured (Fig 6A, S9 Fig). The *sspA* and *pdxA* mutants had significant killing compared to KPPR1. SspA and PdxA were directly linked to oxidative stress as complementation of each restored the ability of *K. pneumoniae* to resist oxidative stress (Fig 6B-C, S9 Fig).

**Fig 6.**
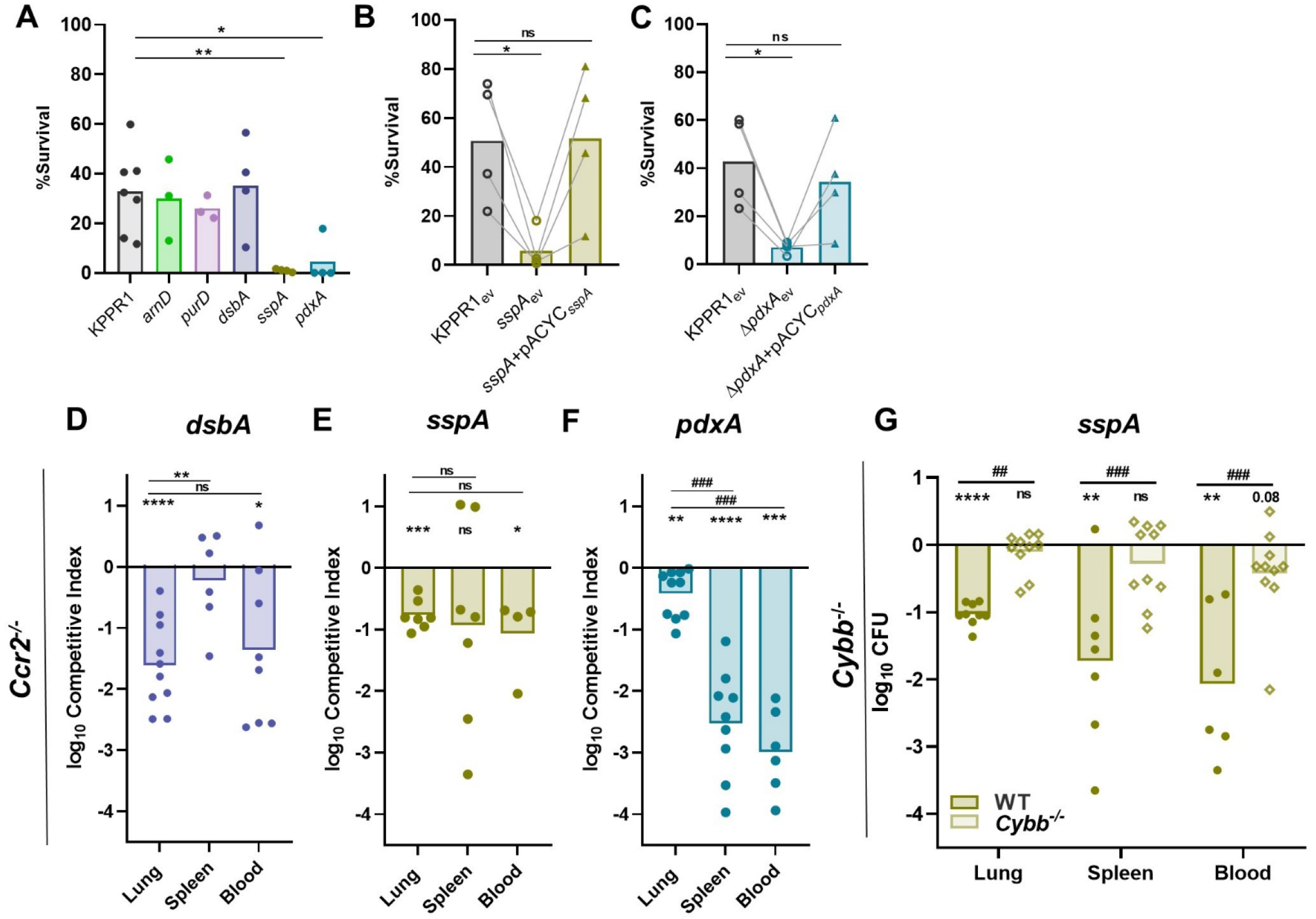
*K. pneumoniae* splenic fitness is maximized by factors relaying resistance to oxidative stress. (A) Resistance to oxidative stress was measured by incubating *K. pneumoniae* strains with H_2_O_2_. Complementation was performed by comparing strains carrying empty pACYC (_ev_) to those with pACYC expression of (B) *sspA* (*sspA*+pACYC_*sspA*_) or (C) *pdxA* (*pdxA*+pACYC_*pdxA*_). (D-G) In a model of bacteremic pneumonia, mice were infected with 1×10^6^ CFU *K. pneumoniae* containing a 1:1 mix of KPPR1 and a transposon mutant for (D) *dsbA*, (E) *sspA*, or (F) *pdxA* in *Ccr*2^*/-*^ mice or (G) *sspA* in *Cybb*^-*/-*^ mice. For (D-G), mean log_10_ competitive index at 24 hours post infection is displayed. In (A-C), *p<0.05, **p<0.01 by one-way ANOVA with Dunnett’s correction for each strain compared to wild-type, n=4. In B-C, lines connect samples from the same replicate. In (D-G), *p<0.05, **p<0.01, ***p<0.001, ****p<0.0001 by one-sample *t* test with a hypothetical value of zero and ^##^p<0.01 by unpaired *t* test. For each group, n≥7 mice in two independent infections.

Inflammatory CCR2^+^ monocytes are recruited to the lung during *K. pneumoniae* infection and are associated with clearance of bacteria and higher rates of murine survival (12, 38, 39). To define if SspA and PdxA enhanced bacteremic pneumonia by interactions with CCR2^+^ monocytes and subsequent oxidative stress, infections were repeated in *Ccr2*^*-/-*^ mice (40). Fitness defects across sites for each strain mirrored that of the wild-type mice and no differences in bacteremia fitness defects were observed between mouse genotypes (Fig 6D-F, S9 Fig). Thus, DsbA, SspA, and PdxA mechanisms enhancing bacteremia fitness likely do not involve interactions with inflammatory monocytes at this early timepoint.

Nox2, phagocyte NADPH oxidase, generates a powerful ROS burst that serves as a main host defense against pathogens. To define if mechanisms mediating oxidative stress resistance were relevant *in vivo*, Nox2 knockout (*Cybb*^*-/-*^) mice were utilized (41). Bacteremic pneumonia was repeated using competitive infections of KPPR1:*sspA* since this mutant showed remarkable susceptibility to oxidative stress and a fitness defect across compartments using this model. In contrast to infections with wild-type mice, SspA was dispensable for fitness across compartments in *Cybb*^*-/-*^ mice (Fig 6G, S9 Fig). Thus, SspA directly increases *K. pneumoniae* bacteremia fitness by promoting resistance to Nox2-mediated oxidative stress in the lung and spleen. Additionally, KPPR1 abundance is significantly increased in the lung of *Cybb*^*-/-*^ mice compared to wild-type mice, demonstrating that Nox2-mediated oxidative stress can partially control abundance of *K. pneumoniae* in the lung (S9 Fig). These data directly link together a *K. pneumoniae* bacteremia fitness factor to a host mechanism of clearance.

## Discussion

In this study, we combined murine intravascular bacteremia and TnSeq technology to define bacterial factors required for *K. pneumoniae* fitness during bloodstream infection. We found that *K. pneumoniae* bacteremia is enhanced by diverse factors, indicating that multiple mechanisms of pathogenesis are deployed to promote the phase of bloodstream survival. Bacteremia fitness factors are distinct in their ability to mediate site-specific fitness, with some genes being required for fitness in one site yet dispensable in others. Replication in serum was supported by purine biosynthesis and endogenous vitamin B6 biosynthesis, emphasizing nutrient restriction in this compartment. The ability of *K. pneumoniae* to resist Nox2-mediated oxidative stress during bacteremia was also critical, and we demonstrate direct interactions between a bacteremia fitness factor and ROS *in vivo*.

Comparing factors required for lung and bloodstream fitness demonstrates that many mechanisms are required across phases of bacteremia, yet others are phase specific. For *K. pneumoniae*, TnSeq has defined a broad spectrum of factors that enhance initial site fitness in the lung (11). *K. pneumoniae* metabolic flexibility is required in the lung through biosynthesis of branched chain (*ilvC/D*) and aromatic (*aroE*) amino acids. However, these factors were not predicted by TnSeq as bloodstream fitness factors in the present study. Other pathways of metabolic flexibility were shared between studies. Multiple members of the purine biosynthesis pathway were predicted to enhance lung and bloodstream fitness, highlighting the importance of this pathway in more than one phase of bacteremia. In contrast, certain factors are unique to later phases of bacteremia. For example, the enzyme GmhB is dispensable for initial site fitness in the lung but was defined by this study and previous work to enhance bloodstream survival (12). Therefore, therapies targeting specific factors may only be relevant in certain phases of disease. By integrating models of direct bacteremia and primary site infection, we were able to study bacteremia fitness factors across distinct phases of pathogenesis.

Infection at primary sites further illuminated contributions of individual factors across phases of bacteremia. For example, DsbA substantially influenced lung fitness while contributing only subtly to splenic fitness across models. SspA and PdxA were also defined as lung fitness factors, and primary site fitness defects increased resolution of splenic defects. It is possible that site-specific stressors in the lung stimulate bacterial defenses that subsequently change splenic fitness. For example, host fatty acid oxidation elicited by other strains of *K. pneumoniae* in the lung results in bacterial adaptation to a new host microenvironment (42). Perhaps these alterations significantly impact fitness at secondary sites, but this has not been experimentally validated for bacteremia. It is also possible that SspA and PdxA contribute to unknown mechanisms of lung dissemination.

Even within the phase of bloodstream survival, different fitness factors are important in different sites of infection. Despite active replication in both organs, *K. pneumoniae* abundance increases in the liver and decreases in the spleen during bacteremia (19, 20). Tissue-resident cells determine differential host responses across sites, and *K. pneumoniae* site-specific fitness has been minimally investigated. Our results demonstrate that multiple *K. pneumoniae* fitness factors contribute to bacteremia through tissue-specific mechanisms as all five factors selected for investigation demonstrated differential fitness patterns across organs (Fig 2). Some factors, like ArnD and DsbA, were required in more than one tissue, while PurD, SspA, and PdxA were required in only one. Thus, the results of this study both illuminate *K. pneumoniae* tissue-specific fitness strategies and define bacterial mutants that can be used as tools to explore host responses at distinct sites.

Gram-negative species actively replicate in the serum, and the biosynthesis of cellular building blocks is critical for survival in the blood since available nutrients differ by site (10, 22). Purine biosynthesis is required for replication in the blood (33), and our TnSeq results demonstrate that many members of the purine biosynthesis operon enhance *K. pneumoniae* bacteremia. Liver fitness was also enhanced by purine biosynthesis, yet splenic fitness did not reach statistical significance for this test. This tissue-specific dynamic for PurD indicates that purine availability may be more restricted in the serum and liver than the spleen. To our knowledge, the source of splenic purines remains unknown and whether the host actively restricts purine availability in the lung, serum, and liver is undefined.

Mechanisms of bacteremia fitness extended beyond metabolic flexibility as TnSeq revealed that many genes supporting splenic fitness were also associated with resistance to oxidative stress. Specifically, we discovered that SspA is required for oxidative stress resistance *in vivo*. Mice lacking the NADPH oxidase Nox2 (*Cybb*^*-/-*^) developed bacteremic pneumonia, yet SspA was dispensable for colonization across sites (Fig 6G) in this background. In contrast, SspA was required for fitness in wild-type mice, which produce a normal ROS response (Fig 5B). This “genetics-squared” approach, comparing infection phenotypes for both *K. pneumoniae* and host mutants allowed resolution of a specific *in vivo* interaction. Thus, the stringent response regulator SspA enhances Gram-negative bacteremia by supporting resistance to oxidative stress, and ROS are responsible for eliciting some portion of *K. pneumoniae* control during bloodstream infection. However, relevant sources of ROS contributing to this response remain unclear. Monocyte-derived macrophages are likely a minimal source of ROS since mice lacking CCR2 do not demonstrate the alleviation of fitness defects in the absence of SspA observed in mice lacking Nox2. Future studies should determine the relevance of neutrophils and other immune cells in this interaction. Notably, SspA displayed tissue-specific fitness during intravascular bacteremia and was required in the spleen yet dispensable in the liver. These differences were not recapitulated *ex vivo* as *sspA* demonstrated no fitness defect in homogenate from either organ. Thus, the oxidative stress either arises from spleen-intrinsic factors that act over a longer time course or from spleen extrinsic factors or cells. In *Francisella tularensis* and *E. coli*, SspA regulates a large network of genes through contact with σ^70^ that alters RNA polymerase transcription of associated genes (43). This dysregulation of a housekeeping sigma factor could create cellular disorder resulting in susceptibility to oxidative and environmental stressors (32). Since the stringent starvation response is a complex system initiated to resist immune cells, redirect metabolism, and promote virulence (44), there could also be multiple additional mechanisms by which SspA supports tissue-specific fitness in addition to ROS resistance.

The use of *ex vivo* organ homogenate did allow resolution of ArnD-mediated tissue-specific fitness. The *arn* operon is well-documented in its role in modifying LPS Lipid A for the repulsion of cationic antimicrobial peptides (31, 45). While ArnD conveyed a significant fitness advantage in the liver and spleen, a surprising finding was that ArnD also conveyed a fitness advantage in the soluble splenic fraction. The fitness defect in the absence of ArnD is likely due to the secretion of an unidentified factor by splenocytes. Together, use of *ex vivo* organ homogenate suggests that factors inherent to individual tissues may inhibit *K. pneumoniae* growth.

A limitations of this study is the investigation of only a small subset of genes revealed as enhancing infection. Remarkably, genes defined by TnSeq as influencing bacteremia both supported and suppressed infection. This study focused on factors encoded by genes in which mutations decreased bacteremia fitness. However, other genes in which mutations led to greater *K. pneumoniae* bacteremia fitness may represent factors that are unfavorable for the bacteria during infection. This subset is also highly diverse and should be mined for future studies further understanding bacteremia fitness dynamics. Another limitation of this study is the ongoing lack of *in vivo* models to study the second phase of bacteremia, dissemination. The ability to measure dissemination separately from lung and splenic fitness would further define relevant interactions for each factor.

Bacteremia is a complex family of infections encompassing multiple sites and responses. Resident cells of the spleen vary widely from those in the liver, and nutrients limited at one site may be abundant in the other. Thus, it is necessary to define factors specifically enhancing bloodstream survival to better understand *K. pneumoniae* bacteremia. This study is the first to define *K. pneumoniae* splenic fitness factors in an *in vivo* mammalian system. Factors enhancing *K. pneumoniae* bacteremia are largely diverse yet represent functions conserved across primary sites and other Gram-negative species and may serve as attractive targets for future therapeutics.

## Materials and Methods

### Murine Bacteremia

This study was performed with careful adherence to humane animal handling guidelines (46) and approved by the University of Michigan Institutional Animal Care and Use Committee (protocol: PRO00009406). Mice used were between 6-12 weeks in age, and each experiment used male and female mice. Wild-type, *Ccr2*^*-/-*^ (40), and *Cybb*^*-/-*^ (41) mice from the C57BL/6 lineage were bred and maintained at the University of Michigan or directly purchased (Jackson Laboratory, Bar Harbor, ME). In each model of bacteremia, *K. pneumoniae* overnight cultures were centrifuged at 5,000xg for 15 minutes and pellets were resuspended in PBS. Cultures were adjusted to the correct concentration based on OD_600_ measurement. To model bacteremic pneumonia and intravascular bacteremia, mice were infected as previously described (12). For pneumonia, animals were anesthetized with isoflurane and a 50µL of PBS containing 1×10^6^ CFU of *K. pneumoniae* was retropharyngeally administered. For intravascular bacteremia, 100µL of PBS containing 1×10^5^ CFU of *K. pneumoniae* was administered by injection via the tail vein. Mice were sacrificed at 24 hours post-inoculation and lung, spleen, liver, or blood were collected. Cardiac punctures were used to obtain whole blood, which was dispensed into heparin coated tubes (BD, Franklin Lakes, NJ). After collection, organs were homogenized in PBS and bacterial burden was calculated by quantitative plating. When appropriate, competitive infections contained a 1:1 ratio of wild-type KPPR1 and a mutant of interest marked with antibiotic selection. Competitive indices were calculated by CFU using the following equation: (*mutant output/wild-type output)/(mutant input/wild-type input)*.

### Bacterial Strains and Reagents

Reagents were sourced from Sigma-Aldrich (St. Louis, MO) unless otherwise noted. *K. pneumoniae* strains were cultured overnight at 37°C with shaking in LB broth (Fisher Bioreagents, Ottowa, ON) or at 30°C on LB agar plates. Media were supplemented with 40µg/mL kanamycin to select for transposon mutants and isogenic knockout strains, or with 50µg/mL chloramphenicol to select for strains containing the plasmid pACYC184 and its derivatives. All bacterial strains in this study are detailed in S3 Table, and primers are detailed in S4 Table.

Complementation plasmids were generated as previously described (12). The complementation vector, pACYC184 was linearized by BamHI and HindIII digestion (New England Biolabs, Ipswitch, MA). The locus for *arnD, purD, sspA, pdxA*, or *pdxA-ksgA-apaG-apaH*, along with upstream regions within 500 base pairs of the open reading frame predicted to contain the native promoter (predicted by SoftBerry BPROM; Softberry Inc, Mount Kisco, NY), were amplified from KPPR1 using primers with 5’ homology to linearized pACYC. For each gene, Gibson assembly was performed using the generated PCR products and linearized pACYC184 according to the manufacturer’s protocol with HiFi DNA Assembly Master Mix (New England Biolabs). The *pdxA*_operon_ amplicon and pACYC184 was ligated after digestion using T4 DNA ligase according to the manufacturer’s protocol (New England Biolabs). The Gibson or ligated product was transformed into *E. coli* TOP10 (New England Biolabs), and constructs were confirmed using full length plasmid sequencing (Plasmidsaurus, Eugene, OR). Verified plasmids were mobilized into *K. pneumoniae* by electroporation.

To generate a *pdxA* (VK055_2525) isogenic knockout, Lambda Red mutagenesis was performed as described (10, 11, 12, 47). Briefly, electrocompetent KPPR1 harboring the pKD46 plasmid was generated using an overnight culture grown at 30°C. The culture was diluted into LB broth with 50µg/mL spectinomycin, 50mM L-arabinose, 0.5mM EDTA (Promega, Madison, WI), and 10µM salicylic acid and grown at 30°C until exponential phase. Cells were cooled on ice for 30 minutes and then pelleted at 8,000xg for 15 minutes. Serial washes were performed at 4°C using 50mL 1mM HEPEs pH 7.4 (Gibco, Grand Island, NY), 50mL diH_2_O, and 20mL 10% glycerol. To generate site specific targets for *pdxA*, a kanamycin resistance cassette from the pKD4 plasmid was amplified with primers also containing 65 base pairs of homology at the 5’ end to the chromosome flanking the *pdxA* open reading frame (S4 Table). This fragment was electroporated into competent KPPR1-pKD46 and transformants were recovered overnight at 30°C, then selected on agar containing kanamycin after a 37° incubation. Knockouts were confirmed with colony PCR using primers flanking and internal to *pdxA*.

### Transposon insertion-site sequencing (TnSeq)

A previously described KPPR1 transposon library consisting of ~25,000 unique random insertions was used to generate four input pools (11, 28). To generate pools, the library was thawed, mixed, and 1mL was removed, pelleted, and resuspended in fresh PBS (Corning, Corning, NY). The OD_600_ was measured, and the library concentration adjusted to 4×10^3^ CFU/mL. 100uL of the adjusted library was plated to achieve a density of ~400 distinct colonies on individual plates. To achieve a desired complexity of 8,500 transposon mutants/pool (S2 Fig), 22 plates were scraped and the CFU combined into PBS. This was repeated four times to generate unique pools (Pools A-D) which were stored at −80°C until use.

Mice were inoculated with one of the four input pools at a dose of 1×10^6^ CFU/mouse in a volume of 100µL via tail vein injection (29). After 24 hours, the mice were euthanized, and spleens were removed and homogenized in 2mL of PBS. Spleen colonization was determined based on quantitative culture using 100µL of homogenate. For each mouse the remaining homogenate was plated in 125µL increments on 100×15mm petri dishes (Corning), incubated at 37°C overnight, scraped, combined in ~125mL PBS, and mixed until homogenous. The OD_600_ was measured and 1×10^9^ CFU was removed, pelleted at 5,000xg for 15 minutes, supernatant removed, and the pellet stored at −80°C until DNA extraction. Any spleen with <8.5×10^3^ CFU/spleen was removed from the study as this colonization is lower than the inoculum mutant complexity and therefore may yield unreliable results. For each input and spleens with appropriate colonization, DNA was extracted from pellets using the Qiagen DNeasy UltraClean Microbial Kit (Qiagen, Hilden DE) according to the manufacturer’s instructions. Purified DNA was submitted to the University of Minnesota Genomics Center for quality verification, library preparation, and sequencing (48). Samples were sequenced using paired-end mode on a NovaSeq 6000 with a depth of 12 million reads/sample. Reads were mapped and normalized as previously described (49), and genes that influence bacteremia fitness were identified using the TnSeqDiff pipeline (50).

### Growth Curves

To assess growth of *K. pneumoniae*, overnights cultures were adjusted to 1×10^7^ CFU/mL in the indicated medium. Using an Eon microplate reader and Gen5 software (Version 2.0, BioTek, Winooski, VT), OD_600_ was measured every 15 minutes and samples were incubated at 37°C with aeration for the duration of the experiment. Strains were measured in the following conditions: LB, M9 salts+20% active human serum, M9 salts+20% heat inactivated human serum, M9 salts+10% murine serum. Differences in growth were detected by area under the curve (AUC, GraphPad Prism Software, LaJolla, CA).

### Hypermucoviscosity

To measure hypermucoviscosity, 500µL of a 1.5mL LB overnight culture for each strain was added to 1.5mL fresh PBS. 900µL of the suspension was used to measure the OD_600_ (pre-spin) while the remaining suspension was centrifuged at 1,000xg for 5 minutes. The OD_600_ of the upper 900µL of supernatant was then measured (post-spin). Hypermucoviscosity=(post-spin)/(pre-spin).

### *Ex vivo* Competition Assays

Uninfected murine spleen and liver were homogenized in 2mL PBS and used for *ex vivo* competition assays as previously described (12). Briefly, 90µL of homogenate and 10µL of PBS containing 1×10^4^ CFU of a 1:1 mixture of *K. pneumoniae* strains were combined and incubated at 37°C for three hours. CFU of each strain at t=0 and t=3 was measured by serial dilutions and quantitative culture from which competitive indices were generated.

### Oxidative Stress Survival Assay

To define *K. pneumoniae* survival in the presence of oxidative stress, overnight bacterial cultures were adjusted to 1×10^7^ CFU/mL in PBS+1mM H_2_O_2_ and incubated for 2 hours at 37°C. Serial dilutions and quantitative culture defined the abundance of each strain before (t=0) and after (t=2) incubation. Percent survival was defined as [(CFU at t=2)/(CFU at t=0)]*100.

### Statistical Analysis

All *in vivo* experiments were performed using at least two independent infections. All *in vitro* and *ex vivo* experiments were performed as independent biological replicates. Statistical significance was defined as a *p-*value <0.05 (GraphPad) as determined using: one-sample tests to assess differences from a hypothetical value of zero for competitive indices, unpaired and paired *t* tests to assess differences between two groups, or ANOVAs with Dunnett’s multiple comparisons to assess differences among multiple groups.

## Acknowledgements

The authors also thank: Dr. Bethany Moore for supplying the *Ccr2*^*-/-*^ mouse lineage; Dr. Gabriel Nunez for supplying the *Cybb*^*-/-*^ mouse lineage; Dr. Jay Vornhagen for assistance with data visualization.

## Supporting Information

**S1 Fig.**
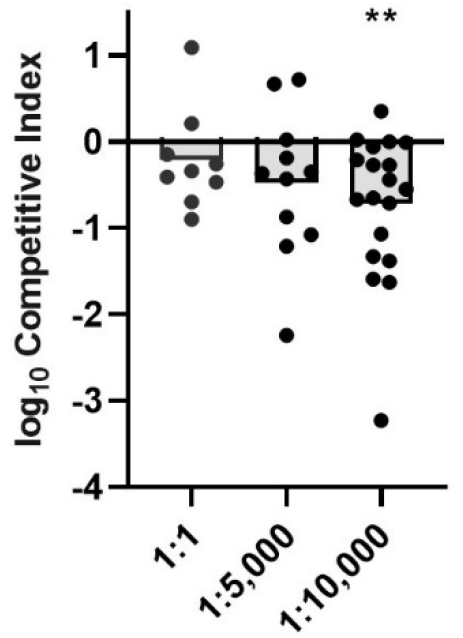
Estimations of experimental bottlenecks in the tail-vein injection model of *K. pneumoniae* bacteremia. Estimation of *in vivo* bottlenecks were determined by competing KPPR1 against a neutral fitness transposon mutant (VK055_*1912*) at varying ratios in the tail vein injection model. **p<0.01 by one sample *t* test with a hypothetical value of 0.

**S2 Fig.**
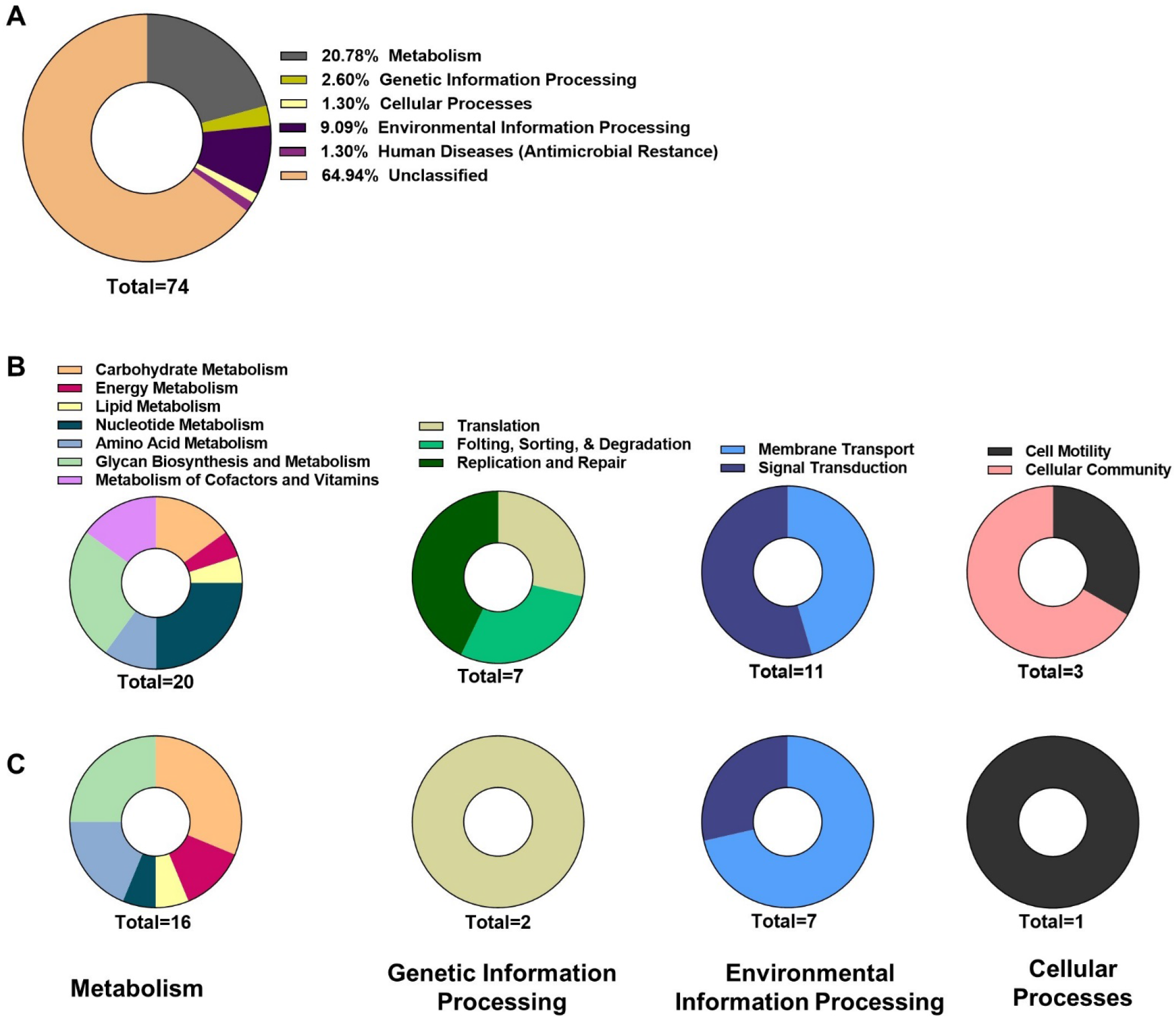
KEGG orthology annotations for genes influencing *K. pneumoniae* bacteremia. (A) Primary KEGG annotations for the 74 genes defined as suppressing fitness. Secondary KEGG annotations for (B) the 58 genes increasing (from Figure 1), or (C) the 74 genes suppressing, *K. pneumoniae* bacteremia fitness. Number=total genes within each annotation, unclassified genes were not included in secondary annotation analysis.

**S3 Fig.**
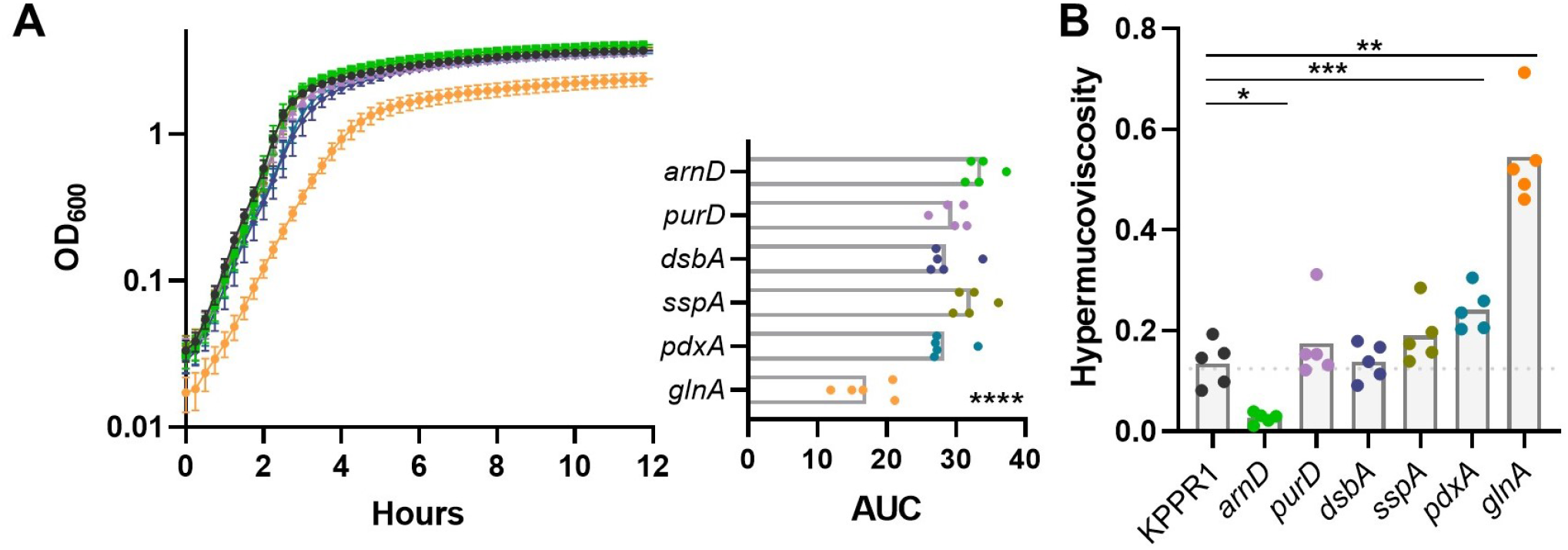
Genes enhancing bacteremia are largely dispensable for *in vitro* replication and have differential effects on hypermucoviscosity. (A) *K. pneumoniae* strains with transposon mutations in genes influencing bacteremia were grown in LB and the OD_600_ was measured every 15 minutes for 12 hours. (B) Hypermucoviscosity was measured for each strain; hypermucoviscosity=(post-spin)/(pre-spin). *p<0.05, **p<0.01, ***p<0.001, ****p<0.0001 by one-way ANOVA with Dunnett’s multiple comparison for each strain compared to KPPR1; n=5.

**S4. Fig.**
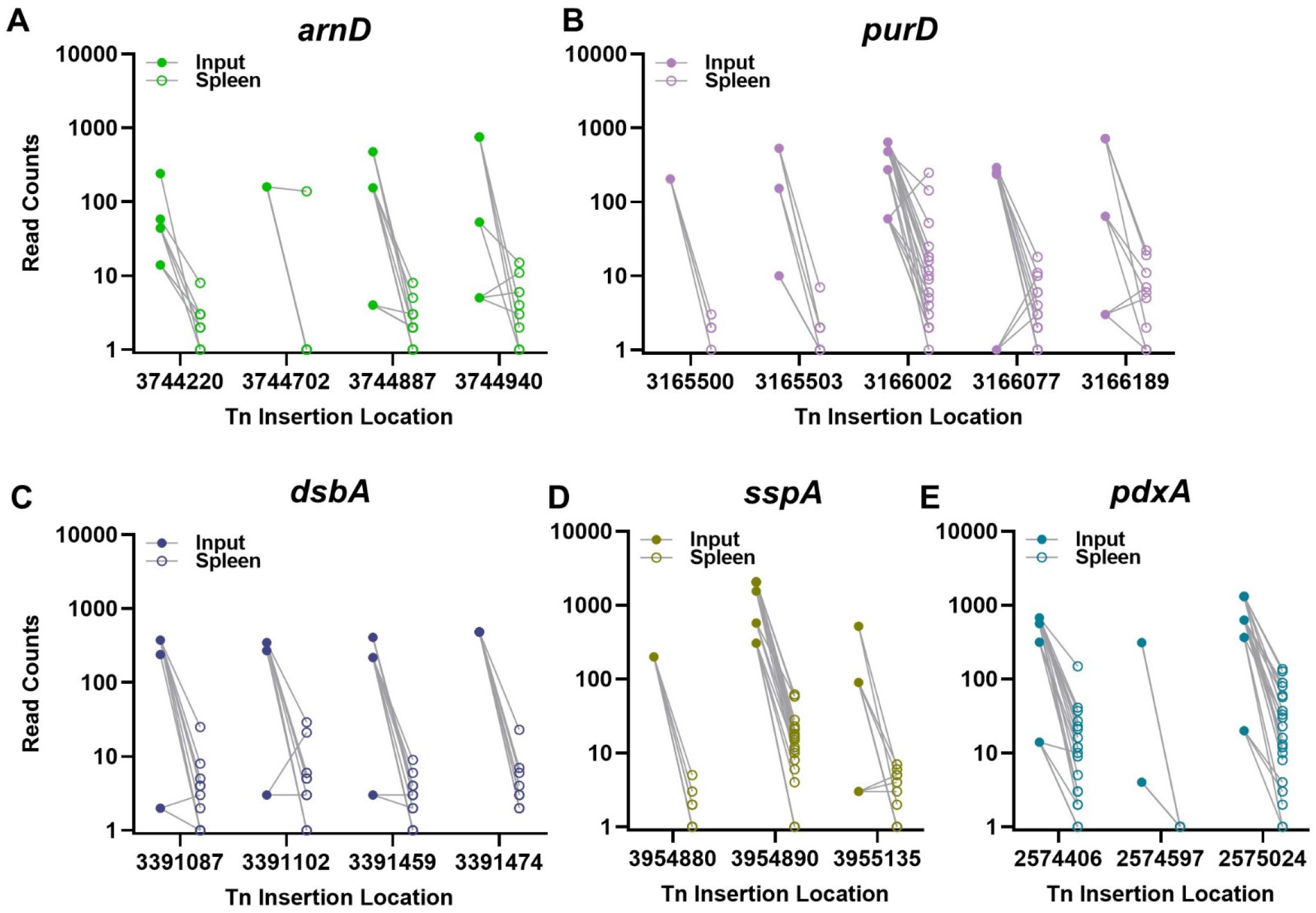
Insertion level analysis for distinct transposon mutations during bacteremia across five fitness factors. Read counts from TnSeq are displayed for unique transposon mutations contained within the genes (A) *arnD*, (B) *purD*, (C) *dsbA*, (D) *sspA*, or (E) *pdxA*. Closed circles indicate input read counts from one of four pools (Pools A-D); open circles indicate read counts from recovered from the spleen 24 hours post infection for individual mice.

**S5 Fig.**
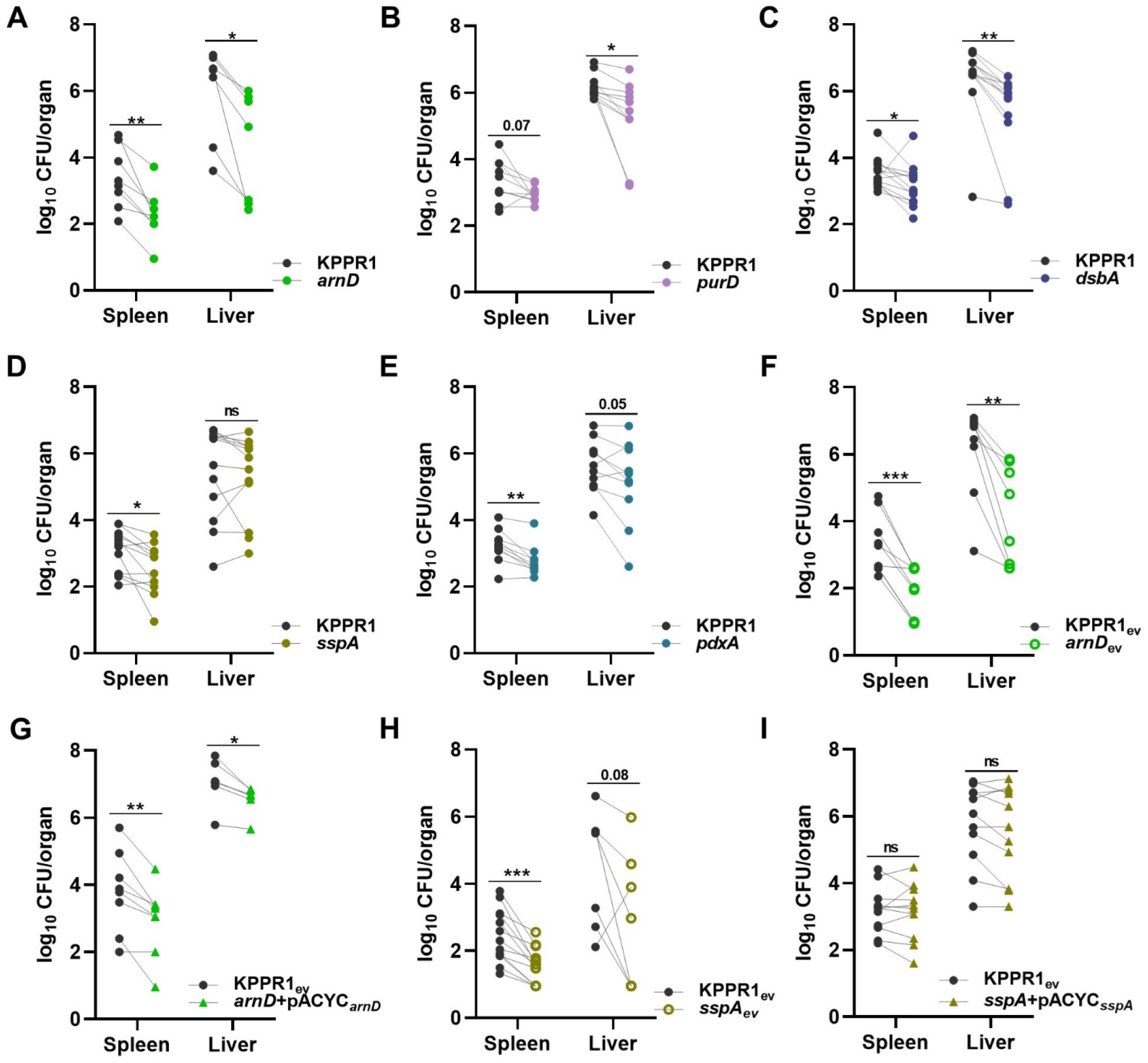
CFU summary for tail vein injections. Five factors indicated by TnSeq as significantly enhancing bacteremia were selected for *in vivo* validation using the tail vein injection model. The 1:1 inoculum consisted of KPPR1 and transposon mutants for (A) *arnD*, (B) *purD*, (C) *dsbA*, (D) *sspA*, or (E) *pdxA*. Competitions were also performed using strains carrying the empty pACYC vector (_ev_) within KPPR1 and (F) *arnD* or (H) *sspA*. Complementation was provided on pACYC under control of the native promoter of (G) *arnD* (*arnD*+pACYC_*arnD*_) or (I) *sspA* (*sspA*+pACYC_*sspA*_). Mean log_10_ CFU burden in the spleen and liver at 24 hours post infection is displayed, corresponding to competitive indices in Figure 2. *p<0.05, **p<0.01, ***p<0.001 by paired t test with Holm-Sidak multiple comparison. For each group, n*≥*8 mice in at least two independent infections.

**S6 Fig.**
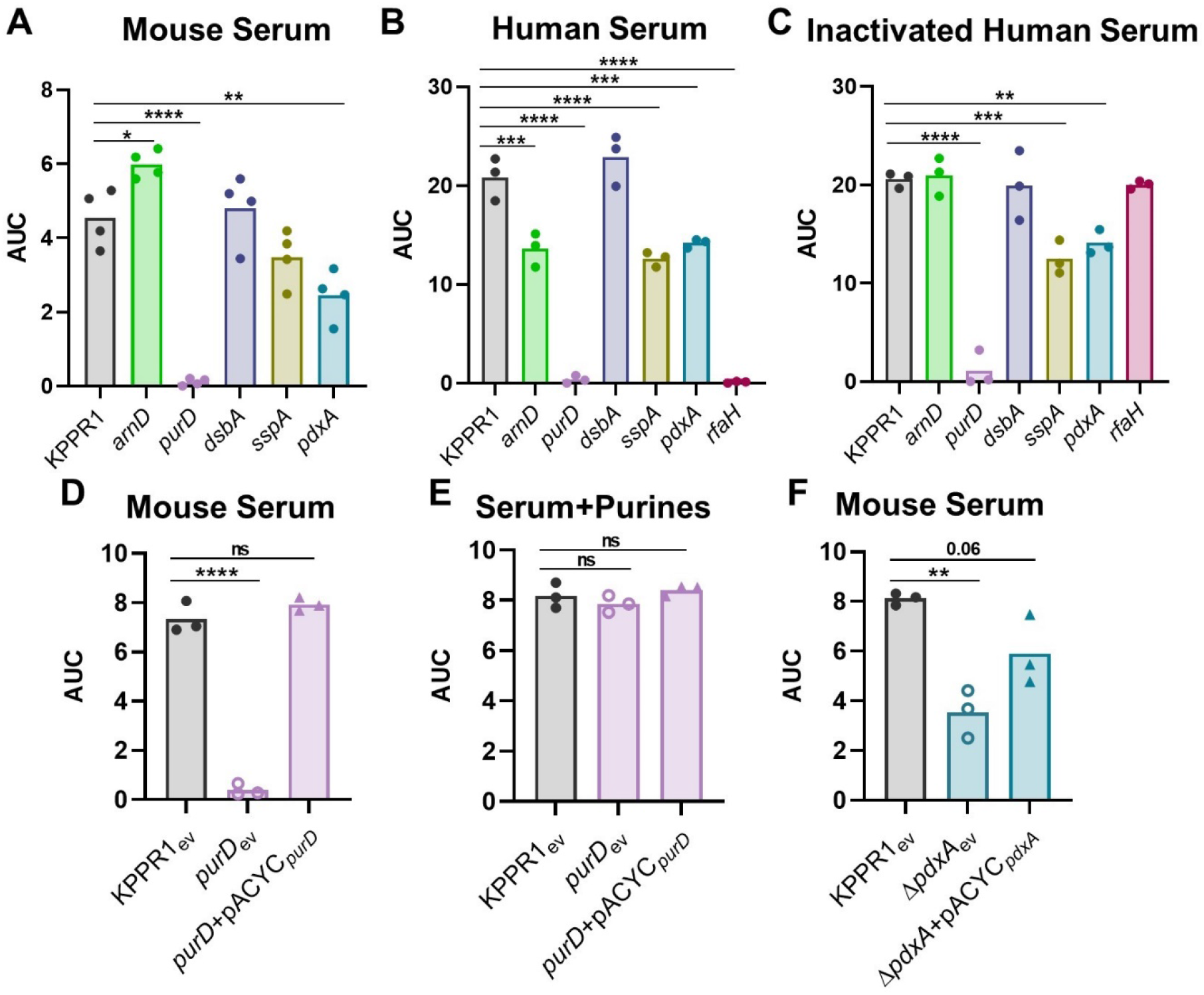
Area under the curve values for *K. pneumoniae* growth in serum. Area under the curve (AUC) was calculated for the growth of individual strains in each condition represented in Figure 3. *K. pneumoniae* strains were grown in M9 salts supplemented with (A) 10% mouse serum or 20% human serum that was either (B) active or (C) heat inactivated. *K. pneumoniae* strains carrying the empty vector pACYC (_ev_) or pACYC expressing (D,E) *purD* (*purD*+pACYC_*purD*_) or (F) *pdxA* (Δ*pdxA*+pACYC_*pdxA*_) were grown in 10% mouse serum (D, F). Chemical complementation for *purD* was measured by supplementation of 1mM purines prior to growth (E). For all, the OD_600_ was measured every 15 minutes for 12 hours. Differences in growth compared to KPPR1 or KPPR1_ev_ were detected by area under the curve using a one-way ANOVA with Dunnett’s multiple comparison for each strain compared to wild-type; *p<0.05, **p<0.01, ***p<0.001, ****p<0.0001. For each, n=3.

**S7 Fig.**
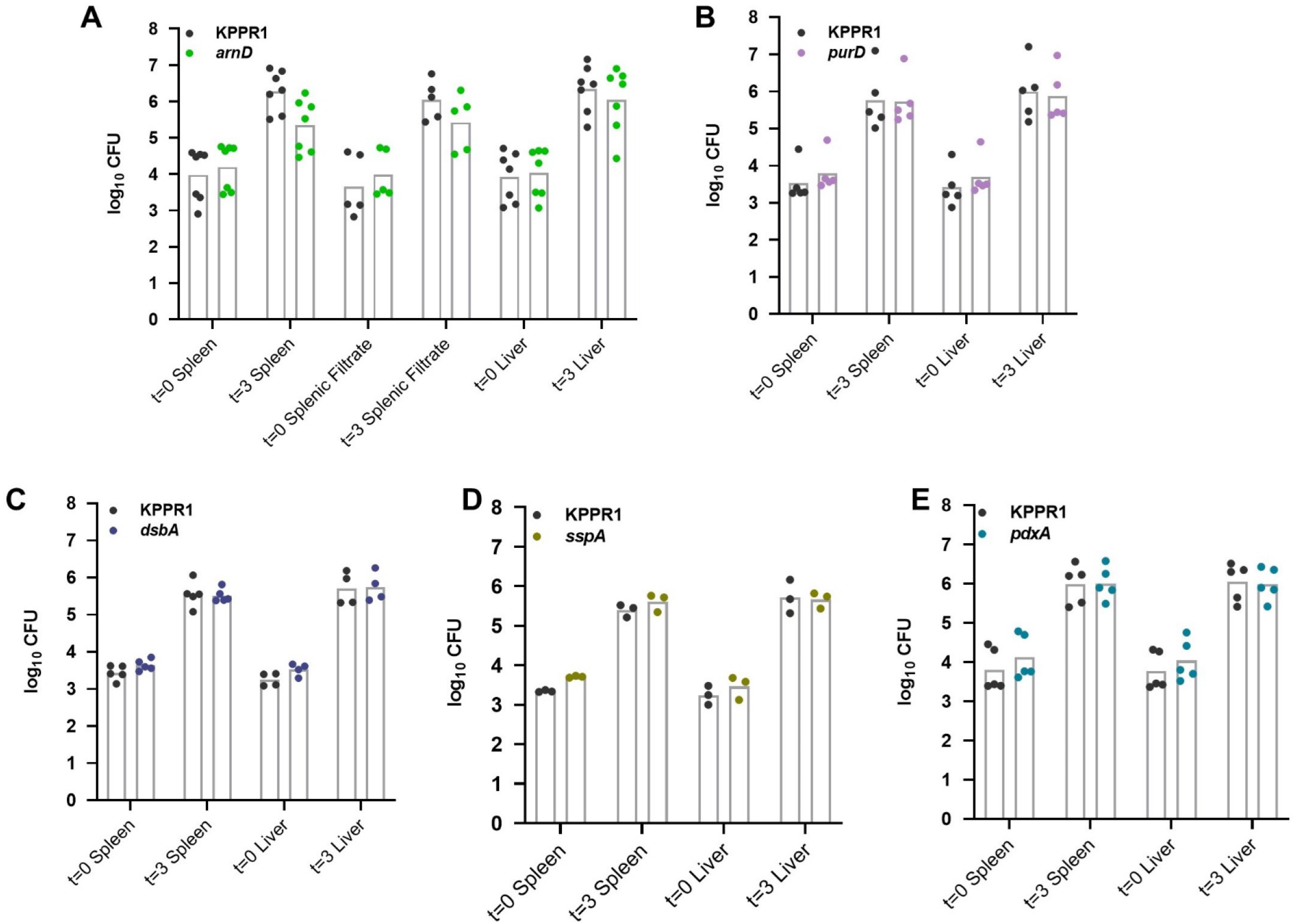
CFU summary for *ex vivo* competitions. *K. pneumoniae* strains were competed at a 1:1 ratio in organ homogenate generated from uninfected mice. An input of 1×10^5^ CFU was added to each well and incubated at 37°C for 3 hours. Log_10_ CFU/well for each strain at the start (t=0) and end (t=3) of the incubation are displayed, corresponding to competitive indices in Figure 4. n≥3 competitions in individual mouse organs.

**S8 Fig.**
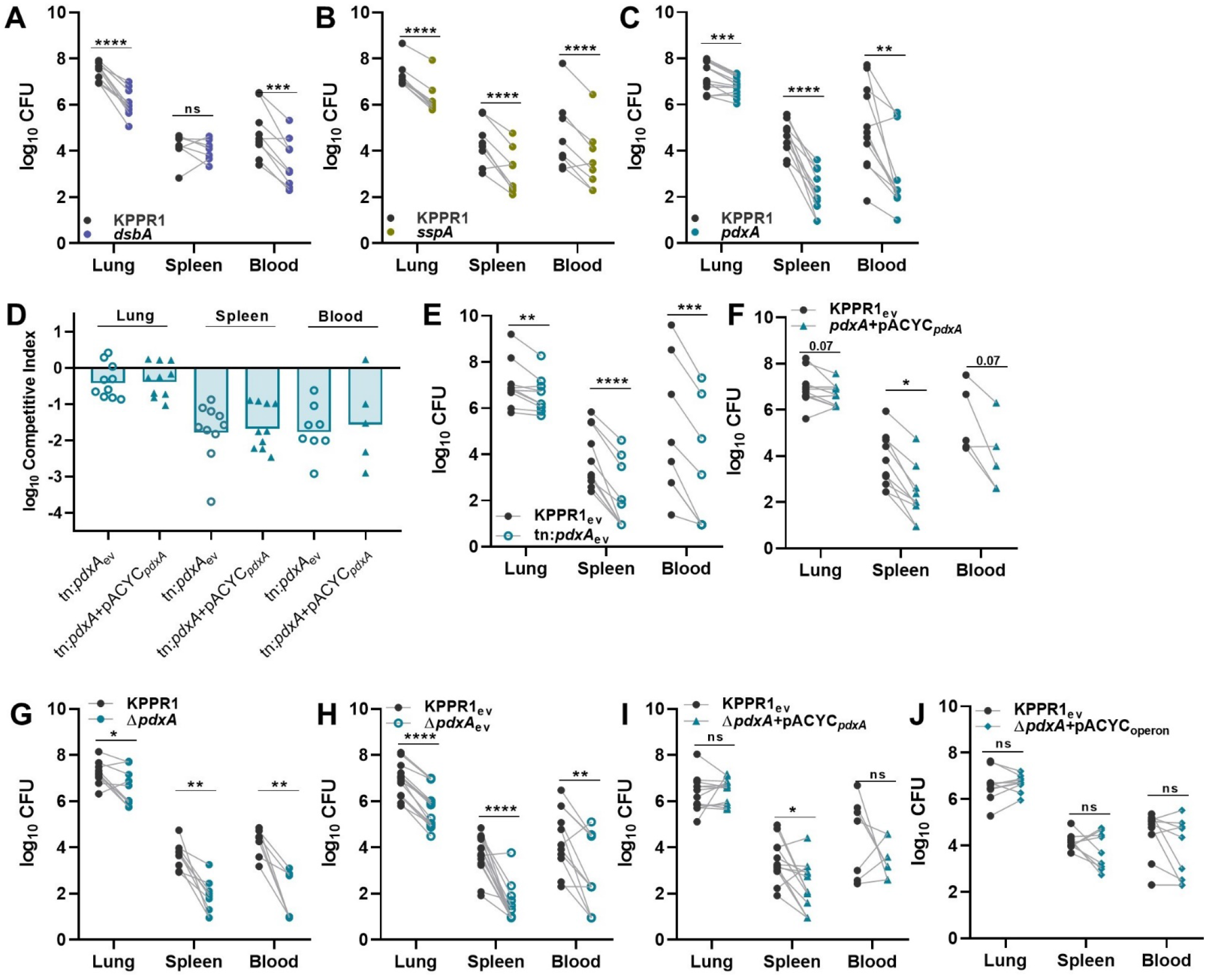
CFU summary for bacteremic pneumonia. To model bacteremic pneumonia, mice were infected with 1×10^6^ CFU *K. pneumoniae*. Competitive infections were performed with a 1:1 mixture of KPPR1 and transposon mutants for (A) *dsbA*, (B) *sspA*, or (C-F) *pdxA* or (G-J) a *pdxA* knockout (Δ*pdxA*). Competitions were performed with strains carrying the pACYC vector (_ev_) or *pdxA* complementation provided on pACYC under control of the native promoter for *pdxA* only (D, F, I; pACYC_*pdxA*_) or *pdxA* and downstream members of the operon (J; pACYC_operon_). Mean log_10_ bacterial burden at 24 hours post infection is displayed corresponding to competitive indices in Figure 5. For (A-C, E-J), *p<0.05, **p<0.01, ***p<0.001, ****p<0.0001 by paired *t* test with Holm-Sidak multiple comparison. For (D) no comparisons were significant between competitive indices within each tissue by unpaired *t* test. For each group, n≥10 mice in at least two independent infections.

**S9 Fig.**
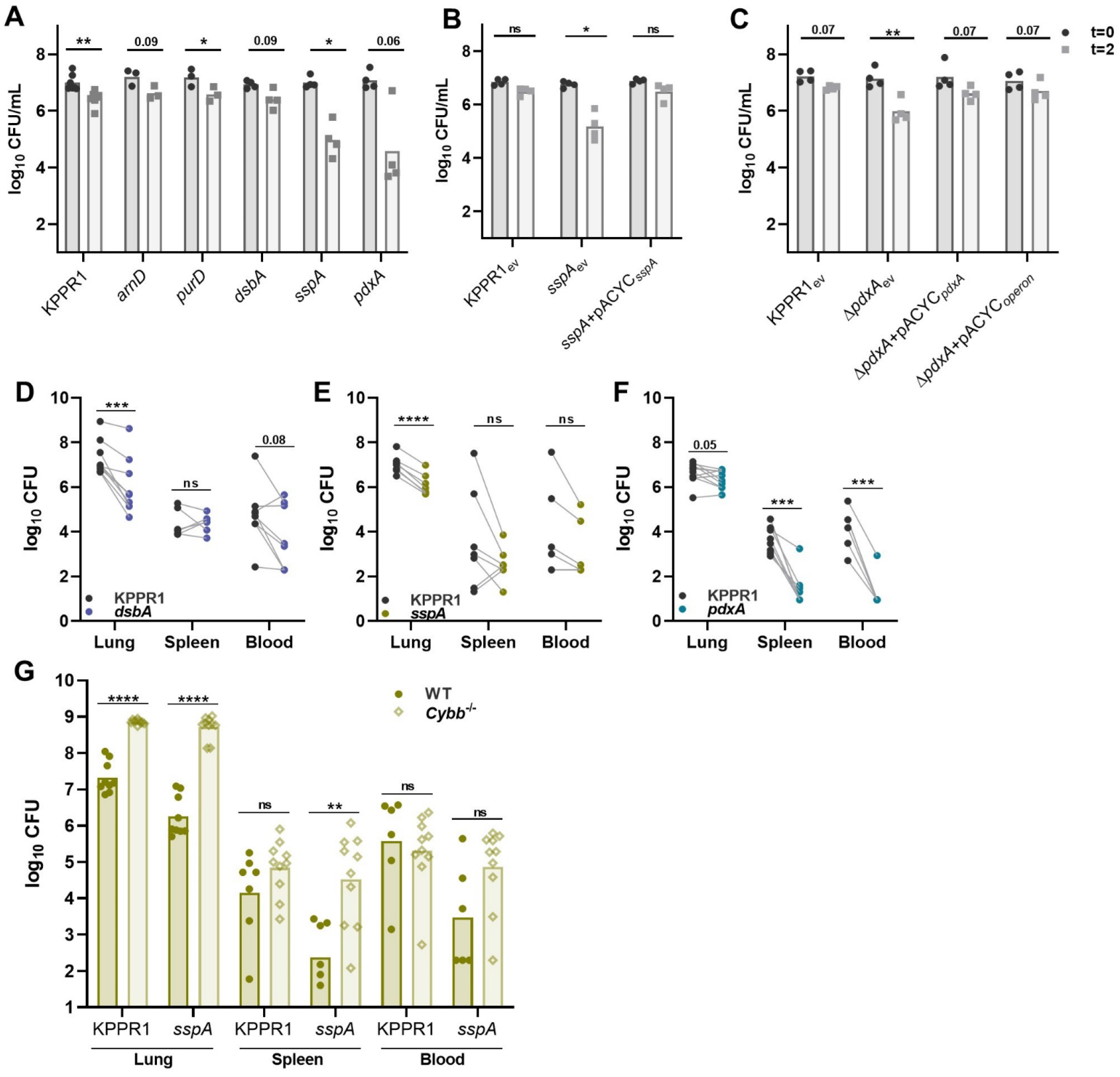
CFU summary for *in vitro* and *in vivo* oxidative stress resistance. (A) Resistance to oxidative stress was measured by incubating *K. pneumoniae* strains with H_2_O_2_. Complementation was performed by comparing strains carrying empty pACYC (_ev_) to those with pACYC expression of (B) *sspA* (*sspA*+pACYC_*sspA*_) or (C) *pdxA* (*pdxA*+pACYC_*pdxA*_). For (A-C), mean log_10_ CFU/mL is displayed. (D-G) In a model of bacteremic pneumonia, mice were infected with 1×10^6^ CFU *K. pneumoniae* containing a 1:1 mix of KPPR1 and a transposon mutant for (D) *dsbA*, (E) *sspA*, or (F) *pdxA* in *Ccr2*^*-/-*^ mice or (G) *sspA* in *Cybb*^-*/-*^ mice. For (A-F), *p<0.05, **p<0.01, ***p<0.001, ****p<0.0001 by paired *t* test with Holm-Sidak multiple comparison. For (G), **p<0.01, ****p<0.0001 by unpaired *t* test. For each group, n≥7 mice in two independent infections.

